# Pericyte and Endothelial Cell Responses within Murine Cerebral Capillaries After Blood Flow Cessation

**DOI:** 10.1101/2025.05.07.652722

**Authors:** Hanaa Abdelazim, Mayd Alsalman, Mason Wheeler, Jessica Pfleger, John C. Chappell

## Abstract

Blood flow provides critical inputs for mechanisms governing vascular homeostasis. Altered hemodynamics can therefore trigger a wide range of cellular responses in blood vessels. Endothelial cells (ECs) downstream of atherosclerotic plaques for instance are exposed to turbulent flow, activating inflammatory pathways that promote immune cell infiltration. In conditions like stroke and myocardial infarction, the abrupt loss of blood flow prompts responses in vascular cells such as ECs and pericytes (PCs) to adapt to ischemic or no-flow conditions. To better understand how cerebral capillary ECs and PCs react to the sudden loss of blood flow, we used a murine brain slice model cultured for 12- and 24-hours in artificial cerebrospinal fluid (aCSF) with 95% oxygen supplementation. As expected, inflammation mediators were upregulated in cultured slices compared to non-cultured samples, particularly those associated with leukocyte recruitment. Additionally, transcriptional markers of extracellular matrix (ECM) remodeling and cell-ECM interactions were elevated, consistent with reduced PC coverage along capillaries. We initially presumed these changes reflected blood-brain barrier (BBB) degradation, but instead we found an increase in mRNA transcripts for EC junctions and stable protein levels for junction molecules, with an apparent rearrangement of Claudin5-based tight junctions. Some capillaries also exhibited reduced diameters, suggesting constriction by PCs or a subset thereof. Consistent with these observations, we found an upregulation of the vasoconstrictor Endothelin-1 (ET-1) with its receptors and contractile proteins found in a subpopulation of PCs. Suppressing ET-1 activity prevented Claudin5 upregulation, indicating that ET-1 might regulate microvascular constriction and associated changes in endothelial tight junctions. Overall, these results suggest that in the absence of blood flow, PCs contribute to capillary wall remodeling by (i) potentially mediating a mechanism driven by ET-1 that affects EC Claudin5 dynamics, and (ii) reducing capillary ECM and detaching from microvessel walls.

## INTRODUCTION

Blood travels throughout an extensive and hierarchical network of vessels, supplying oxygen and nutrients to cells, removing metabolic waste, and distributing hormonal signals across the body. Hemodynamics, or the movement of blood, also generate key biophysical inputs that promote the structural stability of blood vessels^1,2^. For instance, steady, laminar flow promotes the formation and enrichment of capillary endothelial cell (EC) junctions such as Claudin5-based tight junctions^3^. These mechanical forces (e.g. intraluminal pressure) also orchestrate the appropriate synthesis of extracellular matrix (ECM) components that comprise the vascular basement membrane (vBM)^4,5^. Disruption or loss of these mechanical signals can cause damaging changes to the blood vessel walls^6^. Such alterations in blood flow dynamics or the complete cessation of flow can trigger a variety of cellular responses in the affected blood vessels^7^. For example, ECs near a developing atherosclerotic plaque can become exposed to turbulent blood flow, which activates inflammation and promotes the recruitment and infiltration of immune cells^8,9^. Moreover, in conditions like stroke or myocardial infarction, the sudden loss of blood flow leads to several changes within large vessels and the microcirculation, prompting vascular cells to adapt to either a state of no-flow/ischemia or increased blood flow^10^. These responses can have significant effects on vascular health by leading to further complications in the circulatory system.

Across many tissues, and within the brain specifically, additional factors contribute to microvessel integrity. Pericytes (PCs), for example, promote the formation and maintenance of the blood-brain barrier (BBB) through a variety of mechanisms including the induction of EC tight junctions composed of Claudin5^11,12^. PCs also contribute to capillary stability by depositing ECM components into the vBM such as vitronectin, which can be engaged by Integrin-a5 receptors on ECs to enhance vessel integrity^13,14^. Loss of PCs from the blood vessel wall has been implicated in a range of pathologies involving capillary instability and impaired microcirculatory function^15–20^. In particular, a subset of PCs described as “thin-strand PCs” (tsPCs) may be more susceptible to mechanisms leading to their detachment from the microvessel wall^21,22^. To facilitate their outward migration, the ECM surrounding PCs must be remodeled via decreased production of individual components and enzymatic degradation by factors such as matrix metalloproteinases (MMPs)^23–25^. These cellular and molecular contributors to capillary integrity are likely influenced by hemodynamics, as recent studies have begun to describe potential mechanistic connections^26^. But more work is needed to strengthen our understanding of how PCs and the vBM undergo structural rearrangements in response to perfusion changes in microvessel networks.

In addition to detachment and ECM remodeling, PCs can contribute in other ways to capillary adaptations during altered hemodynamics. A PC subpopulation – likely ensheathing PCs (ePCs) and/or mesh PCs (mPCs) – appear to be capable of constricting microvessels and reducing their diameters under certain conditions^27–37^. In particular, contractile PCs have been implicated in narrowing capillary lumens during the sudden loss of flow. Their death while in a constricted state has been suggested to contribute to the “no re-flow” phenomena, that is, limited tissue reperfusion following the successful removal of an upstream blockage^38–41^. Endothelin-1 (ET-1), a potent vasopressor, has been implicated as a driver of PC constriction^42–44^, with a sensitivity to elevated oxygen tension (i.e. hyperoxia)^32^. Interestingly, hypoxia, or abnormally low oxygen levels, as well as disrupted hemodynamics have also been described as stimuli for ET-1 production from ECs^45,46^. Thus, there may be multiple environmental factors that are integrated into mechanisms governing ET-1 release from ECs, indicating a need for further insight into the nuances of ET-1 regulation and its impact on capillary constriction by PC subtypes.

To better understand how ECs and PCs respond to blood flow cessation, particularly within the BBB, here we analyzed microvessels from non-cultured murine brain slices and those cultured in oxygenated artificial cerebrospinal fluid (aCSF) for 12- and 24-hours^47^. In brain slice cultures lacking perfusion, molecular mediators of inflammation, particularly those involved in leukocyte recruitment, were upregulated. Transcriptional markers for ECM remodeling and cell-ECM interactions were also elevated in cortical microvessels, which correlated with reduced pericyte (PC) coverage of cerebral capillaries. These changes suggested BBB breakdown and endothelial degradation, yet mRNA levels for EC junctions increased after 24-hours, with protein levels of junction molecules remaining stable and Claudin5 tight junctions appearing to be structurally rearranged. Additionally, reduced capillary diameters were observed, suggesting that a fraction of cerebral microvessels were likely constricted by PCs. This was further supported by the observed upregulation of the vasoconstrictor ET-1 within cultured brain sections. Consistent with this interpretation, ET-1 receptors and contractile proteins were detected in a PC subset, implying their role in capillary constriction. Suppressing ET-1 activity prevented upregulation of Claudin5/*Cldn5*, indicating a potential link between ET-1 signaling in PCs and EC Claudin5 regulation, with PC constriction contributing to various aspects of cerebral capillary wall remodeling.

## METHODS AND MATERIALS

### In vivo animal studies

In the current study, we utilized brain tissue samples from C57BL/6 and transgenic mice harboring a gene construct labeling microvascular pericytes as well as other cell types expressing Ng2/Cspg4 (e.g. oligodendrocyte precursors), specifically with the DsRed2 fluorophore (STOCK *Tg(Cspg4-DsRed.T1)1Akik/J*, NG2DsRedBAC, The Jackson Laboratory, Strain #008241)^48^. In total, 150 adult mice were used for this study – a mix of males and females between the ages of 3- and 6-months old, under approval granted by the Institutional Animal Care and Use Committee (IACUC), via protocol #23-167. All animals were purchased from The Jackson Laboratory or generated within our in-house animal colony. All experiments involving animal use were performed following review and approval from the IACUC at Virginia Tech. All experimental protocols were reviewed and approved by Virginia Tech Veterinary Staff and the IACUC. The Virginia Tech NIH/PHS Animal Welfare Assurance Number is A-3208-01 (Expires: 31 July 2025). All methods were performed in accordance with the relevant guidelines and regulations provided by these entities. This study was conducted, and is reported, in accordance with ARRIVE guidelines.

### Murine Brain Slice Culture

For each brain slice culture experiment, fresh artificial Cerebrospinal Fluid (aCSF) solutions were prepared. These solutions included a “holding aCSF” formulation prepared as follows: a 10X stock solution of holding aCSF was made containing NaCl (73.63 g/L), KCl (2.24 g/L), NaH₂PO₄ (1.5 g/L), and NaHCO₃ (21.84 g/L). To utilize this solution for an experiment, 100 mL of the 10X stock was diluted with 900 mL of deionized water (DI H₂O), and glucose (1.8 g/L), CaCl₂ (0.5 mL of 1M), and MgSO₄ (4 mL of 1M) were added. The final osmolarity was adjusted to 300-310 mOsm. A “slicing aCSF” solution was also prepared which included the addition of sucrose in the aCSF formulation. This “slicing aCSF” was prepared as follows: a 10X stock solution of slicing aCSF was prepared containing KCl (1.86 g/L), NaH₂PO₄ (1.5 g/L), CaCl₂ (0.74 g/L), and MgSO₄ (24.65 g/L). For experimental use, 100 mL of this stock solution was diluted with 900 mL of DI H₂O, followed by the addition of sucrose (78.73 g/L), NaHCO₃ (2.18 g/L), and glucose (1.8 g/L), adjusting the osmolarity to 315-325 mOsm.

For murine brain extraction, sectioning, slice incubation, and storage, the following methods were utilized. Both male and female adult mice (WT or NG2/*Cspg4^DsRed/+^*) between 3-6 months old were euthanized by isoflurane overdose and rapidly perfused via the heart with ice-cold, oxygenated sucrose-based artificial cerebrospinal fluid (aCSF) i.e. slicing aCSF. The brain was quickly extracted, placed in a sectioning block, and secured onto a vibratome stage. Sucrose-based slicing aCSF and holding aCSF were prepared just before animal euthanasia and oxygenated via bubbling with carbogen (i.e. 95% O₂ and 5% CO₂) and maintained at appropriate temperatures. Sufficient tissue perfusion and exsanguination was confirmed by a reduction in the reddish hue of peripheral organs such as the liver. Extracted brains were submerged in ice-cold sucrose aCSF (slicing aCSF) for uniform cooling. The cerebellum, brainstem, and olfactory bulb were removed before affixing the brain to the vibratome stage while remaining submerged in ice-cold, oxygenated sucrose-based aCSF. Coronal sections were obtained at a thickness of 300 µm using a Leica VT1200S vibratome with continuous carbogen infusion of the aCSF solution. Slices were immediately transferred to a holding chamber containing oxygenated aCSF at room temperature (∼25°C) for recovery. Non-cultured brain slices (i.e. 0-hours in culture) were immediately processed for cortex isolation and downstream analysis. Cultured brain slices were maintained in oxygenated holding aCSF at 32°C for the entire length of incubation period (i.e. 12- or 24-hours) and then processed for cortex isolation and downstream applications. All solutions used herein were prepared to maintain physiological osmolarity, and all procedures were conducted in accordance with established protocols for optimal tissue viability.

### Microvessel Enrichment

Microvascular isolation was performed as described previously^49^. Briefly, enrichment solutions were prepared on ice: Solution 1—HEPES-buffered HBSS, Solution 2—1.8% dextran added to Solution 1, and Solution 3—1% BSA added to Solution 1. Following brain slice generation and incubation, cortices were dissected from the slices. Cortical regions were homogenized in ice-cold Solution 1 using a Dounce tissue grinder and centrifuged (2000×g, 10-min, 4°C). The pellet was resuspended in Solution 2, vortexed, and centrifuged again (4400×g, 15-min, 4°C). The supernatant was removed, and the pellet was resuspended in Solution 3 before filtration through a cell strainer with a 100-µm pore size. The flow-through was collected and centrifuged (4400×g, 5-min, 4°C) and resuspended in Solution 3. Microvascular fractions were collected via an additional centrifugation (2000×g, 5-min, 4°C), and the final pellet was used immediately or stored at -80°C for downstream applications.

### Quantitative RT-PCR

Messenger RNA transcripts were extracted using Trizol, followed by chloroform extraction and isopropanol precipitation. The mRNA pellet was washed with 75% ethanol, air-dried, and resuspended in RNase-free water. mRNA purity and concentration were assessed via a Nanodrop® system. cDNA synthesis was performed using a reverse transcription kit, and qRT-PCR was conducted using TaqMan assays for the probes indicated on an Applied Biosystems QuantStudio6-Flex® platform.

### Western Blot

Protein lysates were prepared by resuspending isolated microvessels in RadioImmunoPrecipitation Assay (RIPA) buffer with protease inhibitors, followed by homogenization and centrifugation (15000×g, 20-min, 4°C). Protein quantification was performed using the Bradford assay, and samples were resolved via SDS-PAGE on 4– 12% Bis-Tris gels. Proteins were transferred to polyvinylidene difluoride (PVDF) membranes and probed with primary antibodies overnight at 4°C: Vitronectin, PECAM-1, Claudin-5, Endothelin-1, α-tubulin, or GAPDH, followed by incubation with secondary antibodies for 2-hrs at room temperature. Bands were detected using a BioRad ChemiDoc® system, ImageLab® software, and the ImageJ/FIJI application.

### Confocal Imaging and Quantitative Image Analysis

Brain slices used for immunostaining and confocal imaging were fixed in 4% paraformaldehyde (PFA) for 30–60-min at room temperature at the corresponding time point of incubation (0-, 12-, and 24-hours) and then washed in PBS. Non-specific antigens within each sample were blocked using PBS-T (0.5% Triton-X 100 in PBS) with 5% bovine serum albumin (BSA) for 1-hr. Samples were then incubated in PBS-T containing primary antibodies overnight at 4°C: PECAM-1, PDGFRβ, Claudin5, ET-1 Type A Receptor (ETA), or aSMA. They were washed in PBS (3 times, 5-min each wash, at room temperature) and incubated with appropriate secondary antibodies for 3-hrs at room temperature, as needed. Nuclei were stained with DAPI (1:1000) for 30-mins at room temperature. Slices were mounted on glass slides with glass coverslips and imaged using a Zeiss LSM 880 confocal microscope with volumetric, z-stack images acquired.

### Quantitative and Statistical Analysis

Statistical analysis was performed using GraphPad Prism 8. For each comparison, a Student t-test or a one-way ANOVA followed by a pairwise Tukey’s post-hoc test was used where applicable. Statistical significance was defined as p ≤ 0.05. Measurements were obtained from a minimum of n = 3 biological replicates per experimental group, with technical replicates included when feasible. Biological replicates were considered to be individual brain samples from a single animal for slices analyzed by confocal imaging. Where the microvessel enrichment protocol was applied, a biological replicate involved the pooling of brain samples from multiple animals such that two (2) brains were required for each biological replicate used for mRNA extraction and qRT-PCR, and four (4) brains were used for each biological replicate for protein analysis by western blot.

## RESULTS

### Inflammation Mediators are Significantly Upregulated in Murine Brain Slices Cultured for 12- and 24-hours in Oxygenated aCSF

Altered hemodynamics and blood flow cessation can induce a broad range of responses in the cellular constituents of affected blood vessels^7^. For example, endothelial cells downstream of a developing atherosclerotic plaque are exposed to turbulent flow, which activates inflammation pathways that lead to immune cell recruitment and infiltration^8,9^. The abrupt loss of blood flow seen in stroke and myocardial infarction also induces vascular cells to respond relative to their new environment i.e. ischemia or no flow vs. heightened perfusion^10^. Here, we sought to better understand how cerebral capillary ECs and PCs respond to the sudden loss of the biophysical inputs from blood flow. To begin addressing this question, we adapted a well-accepted ex vivo murine brain slice model for 12- and 24-hour culture in artificial cerebrospinal fluid (aCSF) infused with 95% oxygen^32,47^. These conditions sustained slice health over the selected experiment durations, with cell death observed by non-viable cell nuclear staining^50^ largely at the cut surfaces and modest levels deeper within these tissues (data not shown). Microvessels were isolated and enriched from cortical regions of cultured brain slices after 12- and 24-hours, allowing us to analyze transcriptional changes within associated cells. We hypothesized that inflammation cues would likely be upregulated, with microvascular ECs responding to altered mechanical inputs by increasing immune cell trafficking mediators. Indeed, we found that transcripts of the proinflammatory cytokine Interleukin-1β (IL1β/*Ilk1b*)^51^ were highly upregulated after 12- and 24-hours in culture relative to non-cultured (i.e. 0-hours in culture) samples (Figure 1A). Similarly, the leukocyte adhesion molecule rapidly expressed by ECs during inflammation onset, E-selectin (*Sele*)^52^, increased in expression significantly at 12- and 24-hours in culture vs. 0-hours (Figure 1B). Vascular Cell Adhesion Molecule-1 (VCAM-1/CD106/*Vcam1*) transcripts significantly increased only at 24-hours in culture (Figure 1C), aligning with its role during the trans-endothelial migration phase of inflammation^53^. Overall, these data indicate that murine brain slices cultured for 12- and 24-hours in oxygenated aCSF undergo an increase in transcriptional signatures of inflammation, consistent with related vascular responses to altered hemodynamics.

**Figure 1.**
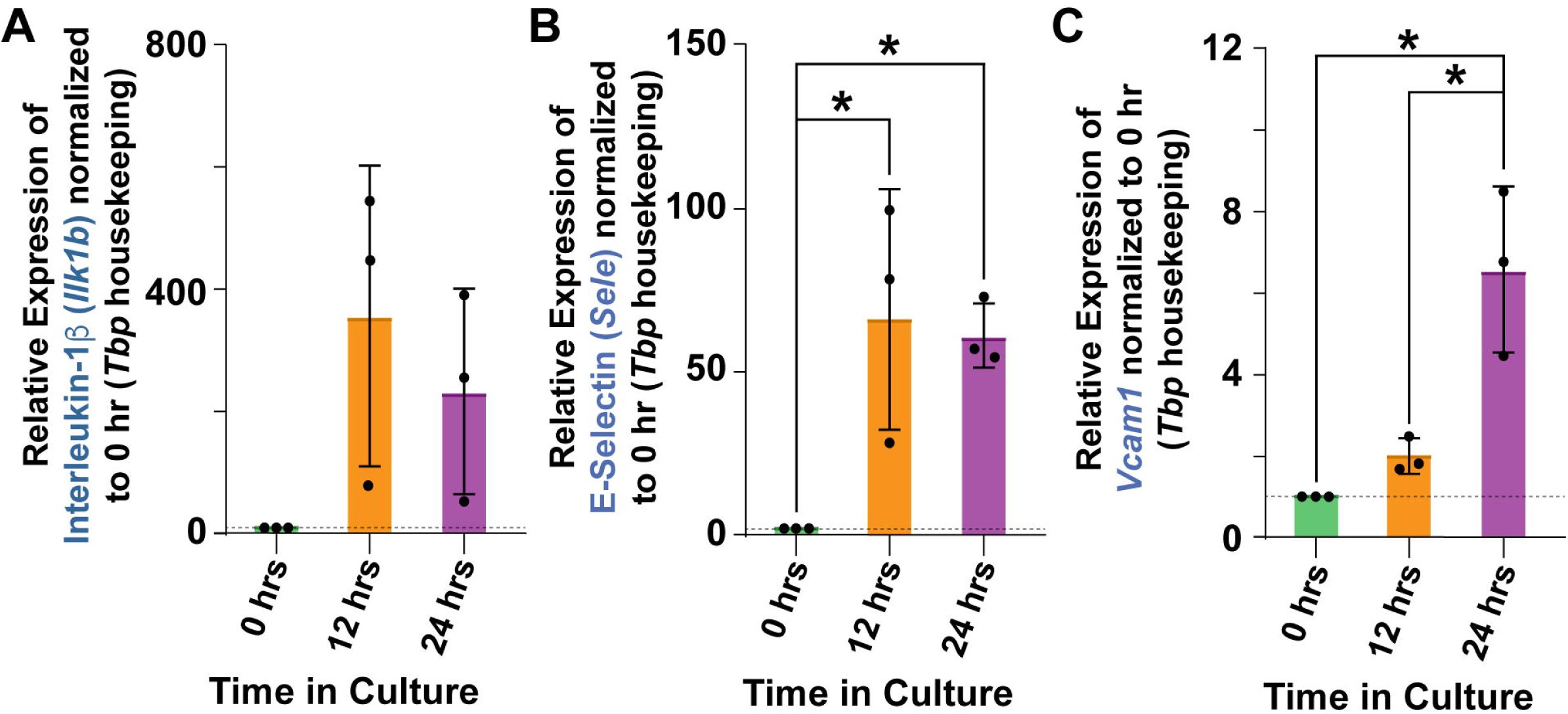
Expression of Immune Cell Recruitment Factors Increases Significantly in Murine Brain Slices Cultured in aCSF for 12- and 24-hours Compared to Non-Cultured Slices. Relative expression of (A) Interleukin-1b (*Ilk-1b*), (B) E-Selectin (*Sele*), and (C) *Vcam1* measured by qRT-PCR in murine brain slices cultured for 12 hours (orange bars) and 24 hours (purple bars), normalized pairwise to 0-hour controls (green bars). *Tbp* served as the housekeeping gene. Filled bars represent averages with error bars as standard deviation. Individual data points are shown for each biological replicate (n=3), and dotted gray lines indicate “1” for referencing to 0-hour control samples. For the comparisons indicated, * denotes p≤0.05 by ANOVA followed by pair-wise Tukey test.

### Increased ECM Remodeling Activity Coincides with Decreased PC Coverage of Cerebral Microvessels in an Ex Vivo Murine Brain Slice Culture Model

Cerebral capillaries, akin to microvasculature in other tissues, are sensitive to sustained changes to their environment including disruptions to the biophysical cues from blood^7,10^. In particular, ECM components can undergo a reduction in their synthesis or targeted degradation by secreted enzymes^23–25^. Capillary PCs also detach from the microvessel wall and migrate outward into interstitial spaces^15–20^. Using the ex vivo murine brain slice model, we asked if these processes occur in a scenario of blood flow cessation but with sufficient oxygen and nutrients available. Matrix metalloproteinase-9 (MMP9/*Mmp9*) has been implicated in eroding the capillary ECM during various disease scenarios^23–25^, and we also found that it was highly upregulated at 12- and 24-hours of brain slice culture compared to non-cultured slices (Figure 2A). Molecules mediating cell attachment to the ECM also displayed transcriptional changes with Integrin-a5 (*Itga5*) in increasing significantly at 24-hours in culture (Figure 2B). Interestingly, transcripts of a glycoprotein within the ECM, vitronectin (*Vtn*)^13,14^, showed a dramatic and significant downregulation at 12- and 24-hours in culture vs. 0-hours slices (Figure 2C).

**Figure 2.**
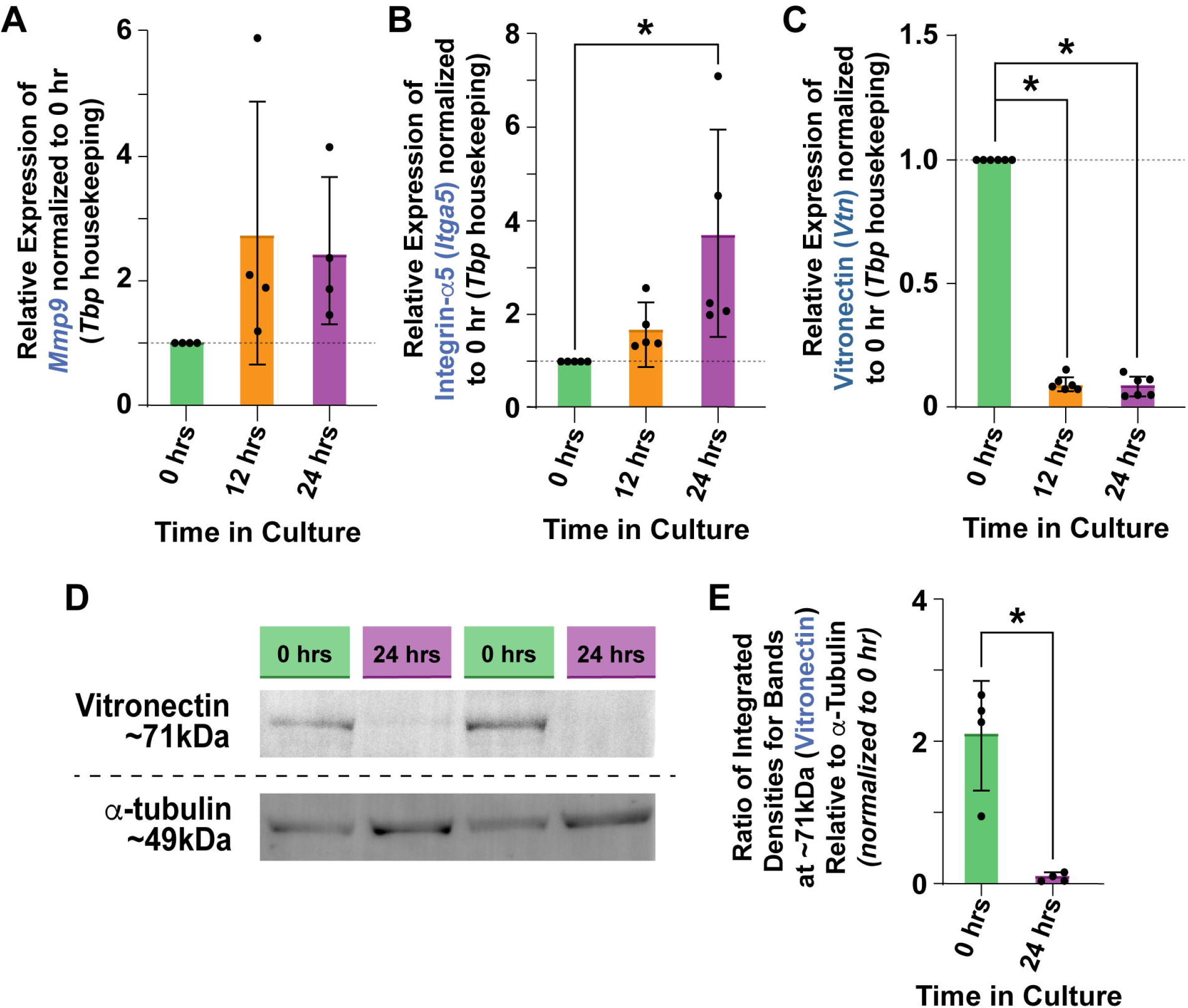
Mediators of ECM Degradation and Engagement are Upregulated in Cultured Brain Slices, while a Key Capillary Wall ECM Component Decreased over 12- and 24-hours. Relative expression of (A) Matrix Metalloproteinase-9 (*Mmp9*), (B) Integrin-a5 (*Itga5*), and (C) Vitronectin (*Vtn*) measured by qRT-PCR in murine brain slices cultured for 12 hours (orange bars) and 24 hours (purple bars), normalized pairwise to 0-hour controls (green bars). *Tbp* served as the housekeeping gene. Filled bars represent averages with error bars as standard deviation. Individual data points are shown for each biological replicate (n>3), and dotted gray lines indicate “1” for referencing to 0-hour control samples. For the comparisons indicated, * denotes p≤0.05 by ANOVA followed by pair-wise Tukey test. (D) Representative western blot images of Vitronectin (∼71kDa) with α-tubulin (∼49kDa) used as a loading control. Four (4) cultured brain slices were pooled for each biological replicate, with n=4 biological replicates per time point (i.e. 0 and 24 hours in culture). *Images were cropped and inverted from original blots, which are provided in Supplemental* Figure 2. (B) Graph of the ratios of integrated densities for bands at the indicated sizes relative to α-tubulin (loading control run on the same blot). Individual data points are shown with averages represented by bars, green for 0-hrs controls and purple for 24 hours in culture. Error bars are standard deviation, and * denotes p≤0.05 by unpaired Student T-test comparison.

Interestingly, Vtn expression in murine brain capillaries remained unchanged in a diet-based model of hyperglycemia (Supplemental Figure 1), suggesting the altered glucose environment during brain slice culture was unlikely driving the decrease in *Vtn* expression. Furthermore, we asked if this notable reduction on the transcriptional level affected vitronectin protein. Western blot analysis revealed a similar decrease in vitronectin content with significantly lower levels at 24-hours in culture relative to non-cultured samples (Figure 2D-E, see Supplemental Figure 2 for original blots), suggesting a unique sensitivity to altered hemodynamics within the cerebral microcirculation.

Observing these ECM-related changes in the brain slice culture model raised several follow-on questions including if a corresponding change in PC coverage was also occurring over these durations. To address this question, we fixed brain slices after 0-, 12-, and 24-hours in culture and used endogenous fluorescent reporters alongside immunostaining to visualize microvascular cells by high-power confocal imaging. Using the NG2*/Cspg4^DsRed/+^* reporter construct^48^, we focused on our analysis on brain regions that included relatively few, if any, NG2/*Cspg4*+ oligodendrocyte precursor cells^54^. As an orthogonal approach, we also applied primary antibodies against (i) full-length Platelet-Derived Growth Factor Receptor-β (PDGFRβ/*Pdgfrb*) to label PCs and (ii) Platelet-Endothelial Cell Adhesion Molecule-1 (PECAM-1/CD31/*Pecam1*) to detect cerebral capillary ECs, with DAPI marking cell nuclei (Figure 3A). In slices cultured for 12- and 24-hours, we observed instances of cells positive for both PC markers in interstitial spaces adjacent to capillary-sized vessels (Figure 3A), suggestive of PC detachment. To gauge the level of PC detachment that may have occurred in this scenario, we measured the percent of DsRed-positive projected area coinciding with PECAM-1-positive projected area. This view of PC coverage revealed a gradual decrease over 12- and 24-hours compared to non-cultured samples (Figure 2B). To complement this measurement, we assessed the ratio of PC soma number per millimeter of vessel length, finding a steady and significant reduction in this ratio at 12- and 24-hours in culture vs. 0-hours (Figure 2C). Collectively, these data suggested that, in murine brain slices cultured in oxygenated aCSF, PC association with cerebral capillaries diminishes over time with corresponding changes in regulators of ECM components that sustain the integrity of the microvascular basement membrane/basal lamina.

**Figure 3.**
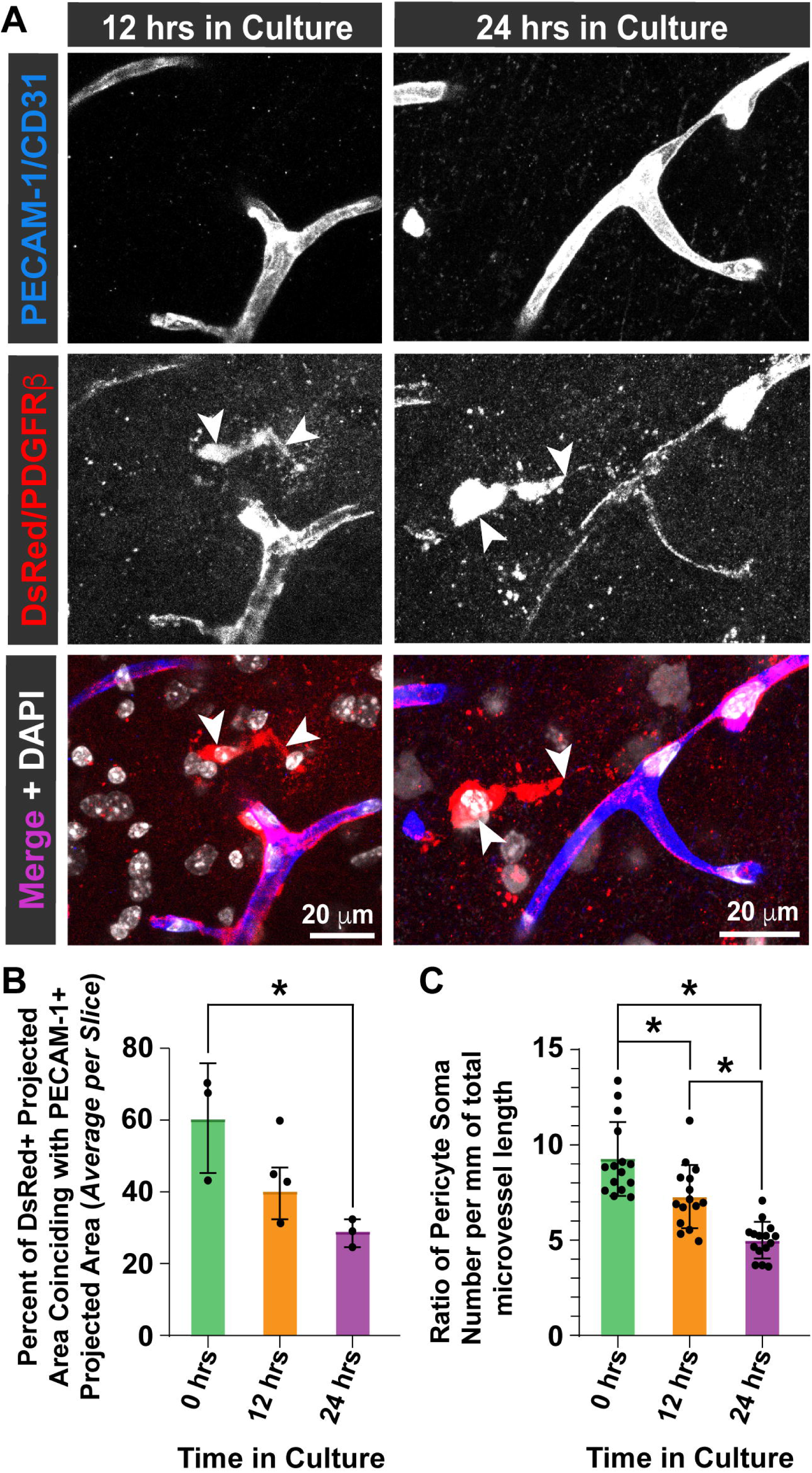
PC Coverage of Cerebral Microvessels Decreased over Time in an Ex Vivo Murine Brain Slice Culture Model. (A) Representative images of NG2/*Cspg4^DsRed/+^* murine brain slices following 12- (i, iii, and v) and 24-hours (ii, iv, and vi) in culture with ECs labeled for PECAM-1/CD31 (i-ii, and blue in v-vi) and PCs labeled by DsRed and PDGFRβ (iii-iv, and red in v-vi). DAPI labeled cell nuclei (white in v-vi). Scale bars, 20 μm. Arrowheads indicate DsRed-/PDGFRβ+ cell soma and extensions in brain interstitial spaces. (B) Graph of the percentage of DsRed-positive projected area coinciding with PECAM-1-positive projected area from slices cultured for 0-hrs (green bar), 12-hrs (orange bar), and 24-hrs (purple bar). Individual data points are shown with bars representing averages from a slice from different animal (n≥3). Error bars are standard deviation, and * denotes p≤0.05 by Student T-test comparison. (C) Graph of the ratio of DsRed-/PDGFRβ-positive cell soma number per millimeter of total PECAM-1-positive microvessel length from slices cultured for 0-hrs (green bar), 12-hrs (orange bar), and 24-hrs (purple bar). Individual data points are shown with bars representing averages from a slice from different animal (n≥3). Error bars are standard deviation, and * denotes p≤0.05 by Student T-test comparison.

### Transcripts for EC Junction Molecules are Upregulated over 24-hours in Murine Brain Sections Exposed to Oxygen-infused aCSF, with Claudin5 Undergoing Morphological Rearrangement

Loss of ECM stability and PC detachment have been associated with disrupted EC junctions, particularly in the context of compromised BBB stability^11,12^. Upon detecting ECM- and PC-related changes within microvessels of the brain slice culture model, we asked if EC junctions also unraveled in the absence of blood flow under these specific culture conditions. PECAM-1 for instance supports EC junctions and has been implicated in stabilizing the BBB, though it also facilitates leukocyte transmigration through the endothelium during inflammation^55^. Therefore, we assessed *Pecam1* transcript levels in microvessels at 12- and 24-hours and found a significant increase at the 24-hour time point relative to 0-hours in culture (Figure 4A). We found no sensitivity to *Pecam1* expression in murine cerebral capillaries within a diet-induced model of hyperglycemia (Supplemental Figure 1), suggesting glucose levels in the slice culture model were not likely causing the observed transcriptional increase. PECAM-1 protein levels at 24-hours displayed some variability, but no significant changes were detected (Figure 4B-C, see Supplemental Figure 3 for original blots). We continued investigating potential alterations in mediators of BBB stability by analyzing the EC tight junction Claudin5 (*Cldn5*), as it has been heavily implicated in this role and has been described as sensitive to ECM and PC perturbations^11,12^. Surprisingly, we found that, similar to *Pecam1* transcripts, *Cldn5* was upregulated significantly at 24-hours in culture compared to non-cultured slices (Figure 4D), but not in a model of diet-induced hyperglycemia (Supplemental Figure 1). Claudin5 protein levels fluctuated at 24-hours, but they were not significantly altered over the 24-hour duration of brain slice culture (Figure 4E-F, see Supplemental Figure 4 for original blots).

**Figure 4.**
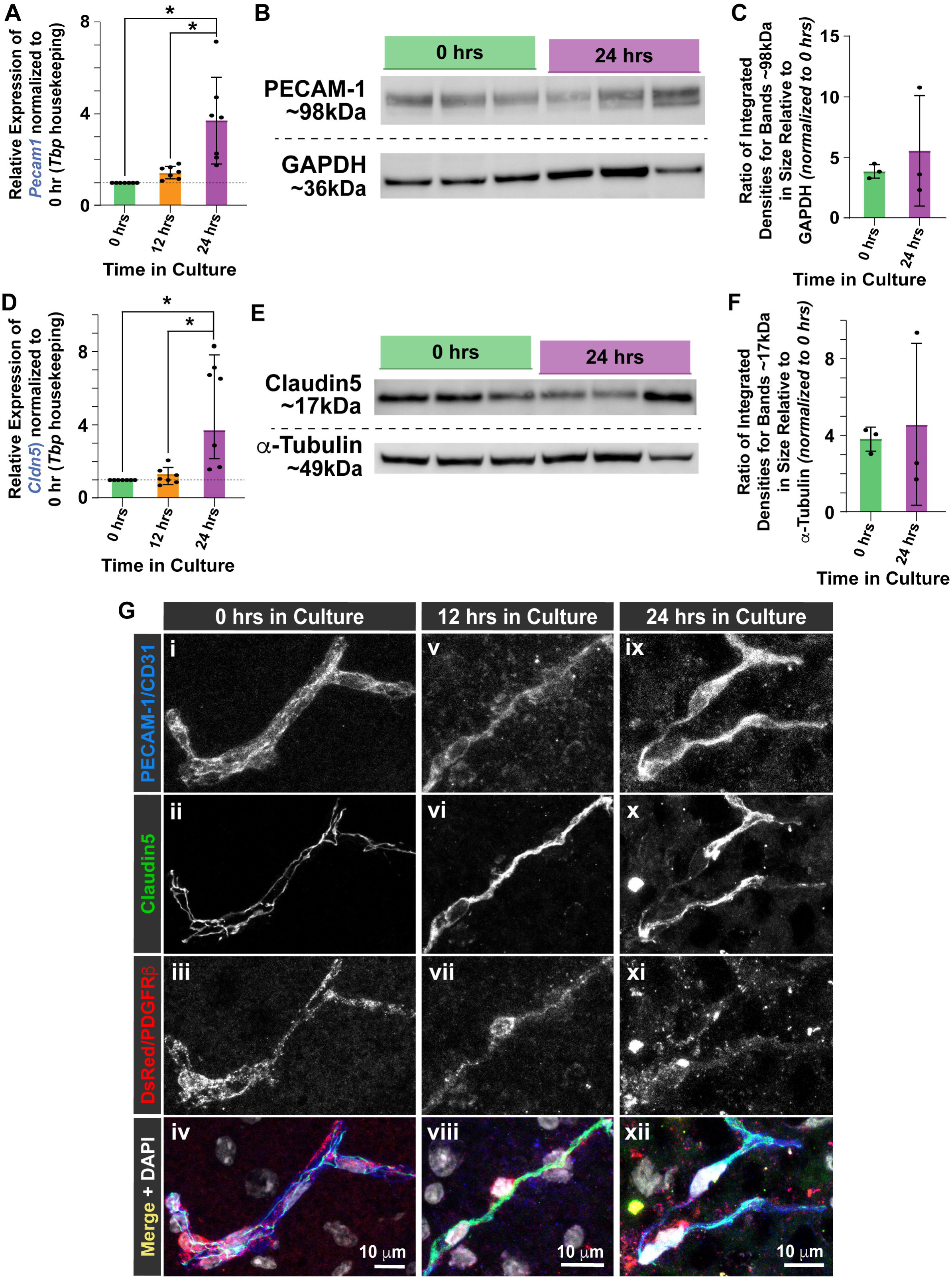
EC Junction mRNA Transcripts are Upregulated over 24-hours in Murine Brain Slices Cultured in Oxygenated aCSF, with Claudin5-based Tight Junctions Undergoing Morphological Rearrangements. (A) Relative expression of *Pecam1* measured by qRT-PCR in murine brain slices cultured for 12 hours (orange bars) and 24 hours (purple bars), normalized pairwise to 0-hour controls (green bars). *Tbp* served as the housekeeping gene. Filled bars represent averages with error bars as standard deviation. Individual data points are shown for each biological replicate (n≥6), and dotted gray lines indicate “1” for referencing to 0-hour control samples. For the comparisons indicated, * denotes p≤0.05 by ANOVA followed by pair-wise Tukey test. (B) Representative western blot images of PECAM-1 (∼98kDa) with GAPDH (∼36kDa) used as a loading control. Four (4) cultured brain slices were pooled for each biological replicate, with n=3 biological replicates per time point (i.e. 0 and 24 hours in culture). *Images were cropped and inverted from original blots, which are provided in Supplemental* Figure 3. (C) Graph of the ratios of integrated densities for bands at the indicated size (∼98kDa) relative to GAPDH (loading control run on the same blot). Individual data points are shown with averages represented by bars, green for 0-hrs controls and purple for 24 hours in culture. Error bars are standard deviation. (D) Relative expression of *Cldn5* measured by qRT-PCR in murine brain slices cultured for 12 hours (orange bars) and 24 hours (purple bars), normalized pairwise to 0-hour controls (green bars). *Tbp* served as the housekeeping gene. Filled bars represent averages with error bars as standard deviation. Individual data points are shown for each biological replicate (n≥6), and dotted gray lines indicate “1” for referencing to 0-hour control samples. For the comparisons indicated, * denotes p≤0.05 by ANOVA followed by pair-wise Tukey test. (E) Representative western blot images of Claudin5 (∼17kDa) with α-tubulin (∼49kDa) used as a loading control. Four (4) cultured brain slices were pooled for each biological replicate, with n=3 biological replicates per time point (i.e. 0 and 24 hours in culture). *Images were cropped and inverted from original blots, which are provided in Supplemental* Figure 4. (F) Graph of the ratios of integrated densities for bands at the indicated size (∼17kDa) relative to α-tubulin (loading control run on the same blot). Individual data points are shown with averages represented by bars, green for 0-hrs controls and purple for 24 hours in culture. Error bars are standard deviation. (G) Representative images of NG2/*Cspg4^DsRed/+^* murine brain slices following 0- (i-iv),12- (v-viii) and 24-hours (ix-xii) in culture with ECs labeled for PECAM-1/CD31 (i, v, and ix, and blue in iv, viii, and xii) and for Claudin5 (ii, vi, and x, and green in iv, viii, xii), and PCs labeled by DsRed and PDGFRβ (iii, vii, and xi, and red in iv, viii, xii). DAPI labeled cell nuclei (white in iv, viii, and xii). Scale bars, 10 μm.

Based on previous studies and commentaries^11,12,56–58^, we had expected to observe decreases in Claudin5 transcripts and protein; however, we observed trends that did not align with this expectation. To better understand this discrepancy, we designed a follow-on analysis to visually observe potential morphological changes in EC junctions. Specifically, we immunostained slices from NG2*/Cspg4^DsRed/+^* animals for PECAM-1 and PDGFRβ as above, adding an additional step of antibody labeling Claudin5 alongside staining nuclei with DAPI (Figure 4G). For non-cultured samples (i.e. 0-hours in culture), we found thin bands of Claudin5 signal aligning with PECAM-1-positive junctions in microvessels associated with cells positive for PC markers. In contrast, capillaries within samples cultured for 12- and 24-hours displayed much thicker Claudin5-positive bands and were within vessels with smaller outer diameters (Figure 4G). We applied an orthogonal view to confocal images taken as a z-stack, to better assess the apparent morphological rearrangement of Claudin5 within these vessels. Interestingly, we found the thin bands of Claudin5 signal located predominantly at the outer edge of cerebral capillaries at 0-hours, whereas the thicker Claudin5 signal in vessels of 24-hour samples appeared to radiate out from the center towards the outer surface (see Supplemental Figure 5). Taken together, these data contrasted with our initial expectation of degraded EC junctions, showing signs more in the direction of transcriptional upregulation and altered morphology with sustained protein levels.

### ET-1 Increased within Murine Brain Slices Cultured for 12- and 24-hours in aCSF

Altered hemodynamics, and specifically lower shear stress, causes a sustained release of the potent vasopressor ET-1 from ECs^46^. This published observation prompted our hypothesis that ET-1 might be upregulated in the murine brain slice culture model given the dramatic shift in the biophysical inputs experienced by cerebral capillaries. We assessed ET-1 (*Edn1*) transcripts from murine brain microvessels collected at 12- and 24-hours in culture and found significant increases compared to non-cultured samples (Figure 5A). Again, altered glucose levels did not appear responsible for this change, as diet-induced hyperglycemia did not affect *Edn1* expression in murine cerebral microvessels (Supplemental Figure 1). ET-1 protein levels at 0- and 24-hours were compared by western blot, and an increase in ET-1 protein was found at the 24-hour time point, though not statistically significant (Figure 5B-C, see Supplemental Figures 6 and 7 for original blots and additional replicates). Thus, while data on the protein level did not show a substantial increase in ET-1 levels in cultured brain slices, we did detect a significant transcriptional upregulation, consistent with previous reports of ET-1 regulation in ECs being sensitive to hemodynamic changes.

**Figure 5.**
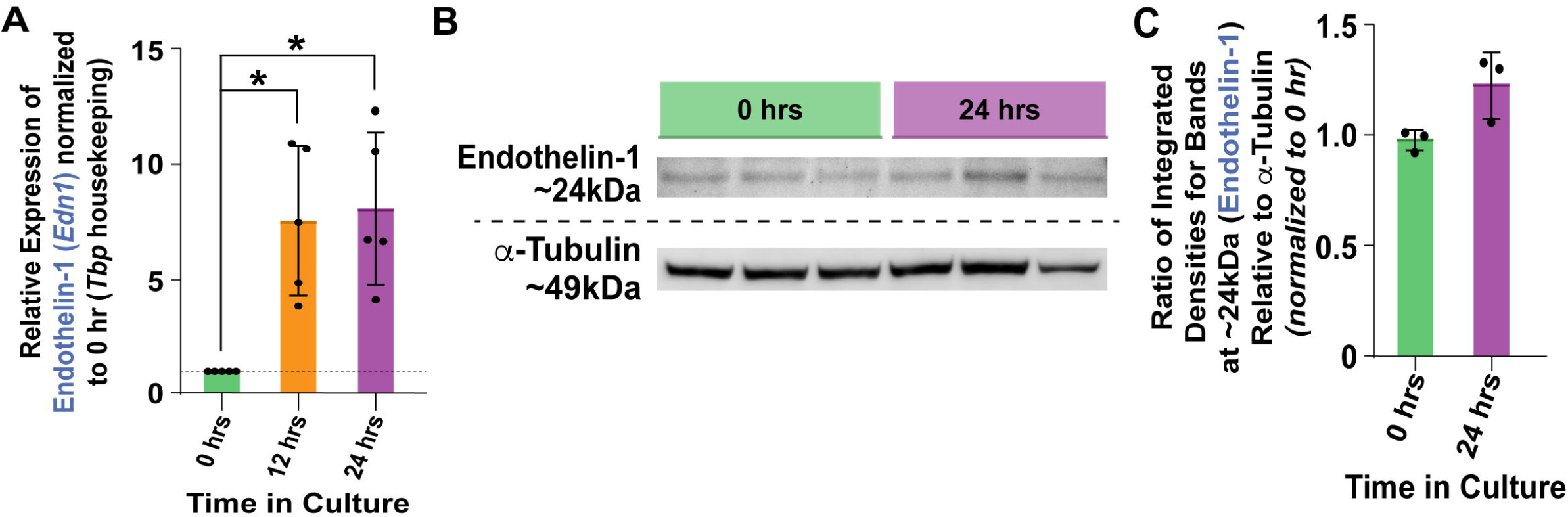
ET-1 is Upregulated in Murine Brain Slices Cultured for 12- and 24-hours in oxygen-bubbled aCSF. (A) Relative expression of *Edn1* measured by qRT-PCR in murine brain slices cultured for 12 hours (orange bars) and 24 hours (purple bars), normalized pairwise to 0-hour controls (green bars). *Tbp* served as the housekeeping gene. Filled bars represent averages with error bars as standard deviation. Individual data points are shown for each biological replicate (n=5), and dotted gray lines indicate “1” for referencing to 0-hour control samples. For the comparisons indicated, * denotes p≤0.05 by ANOVA followed by pair-wise Tukey test. (B) Representative western blot images of Endothelin-1 (∼24kDa) with α-tubulin (∼49kDa) used as a loading control. Four (4) cultured brain slices were pooled for each biological replicate, with n=3 biological replicates per time point (i.e. 0 and 24 hours in culture). *Images were cropped and inverted from original blots, which are provided in Supplemental* Figure 6*. Additional replicates are shown in Supplemental* Figure 7. (C) Graph of the ratios of integrated densities for bands at the indicated size (∼24kDa) relative to α-tubulin (loading control run on the same blot). Individual data points are shown with averages represented by bars, green for 0-hrs controls and purple for 24 hours in culture. Error bars are standard deviation.

### A PC Subset Harbors ET-1 Type A Receptors and **α**SMA expression, and Inhibiting ET-1 Activity over 24-hours Prevented Upregulation of Claudin5 Transcripts

Considering published studies^42–44^ alongside the observations described above, we constructed an overarching hypothesis for testing follow-on questions. Specifically, we asked if cerebral capillary PCs, or perhaps a subset, might be responsive to increased ET-1 levels via cell surface receptors. These signals might in turn stimulate this PC subpopulation to constrict a portion of capillaries and contribute to the observed changes in Claudin5 transcription and morphology (see Supplemental Figure 8 for a summary schematic). To begin dissecting our working model of cerebral PC and EC responses to altered hemodynamics, we immunostained non-cultured murine brain slices from NG2*/Cspg4^DsRed/+^* animals for PECAM-1 and ET-1 Type A Receptors (ETA), with cell nuclei labeled by DAPI (Figure 6A). We found a distinct region of the capillary wall positively labeled for ET-1 receptors that coincided with DsRed signals, with both located directly adjacent to the PECAM-1-positive endothelium. We validated the accuracy of the ETA primary antibody by also imaging larger diameter vessels containing DsRed-positive vascular smooth muscle cells (vSMCs) that are well known to express ET-1 Type A Receptors on their cell surface (see Supplemental Figure 9). Furthermore, we drew from several established single cell RNA-sequencing databases to gather evidence aligning with the presence of *Ednra* (ETA) transcripts in PC populations from various sources (see Supplemental Figure 10)^59–61^. Overall, these data support the idea that PCs, or at least a subset, express ET-1 Type A Receptors that may elicit intracellular signals to orchestrate cell behaviors such as contractility.

**Figure 6.**
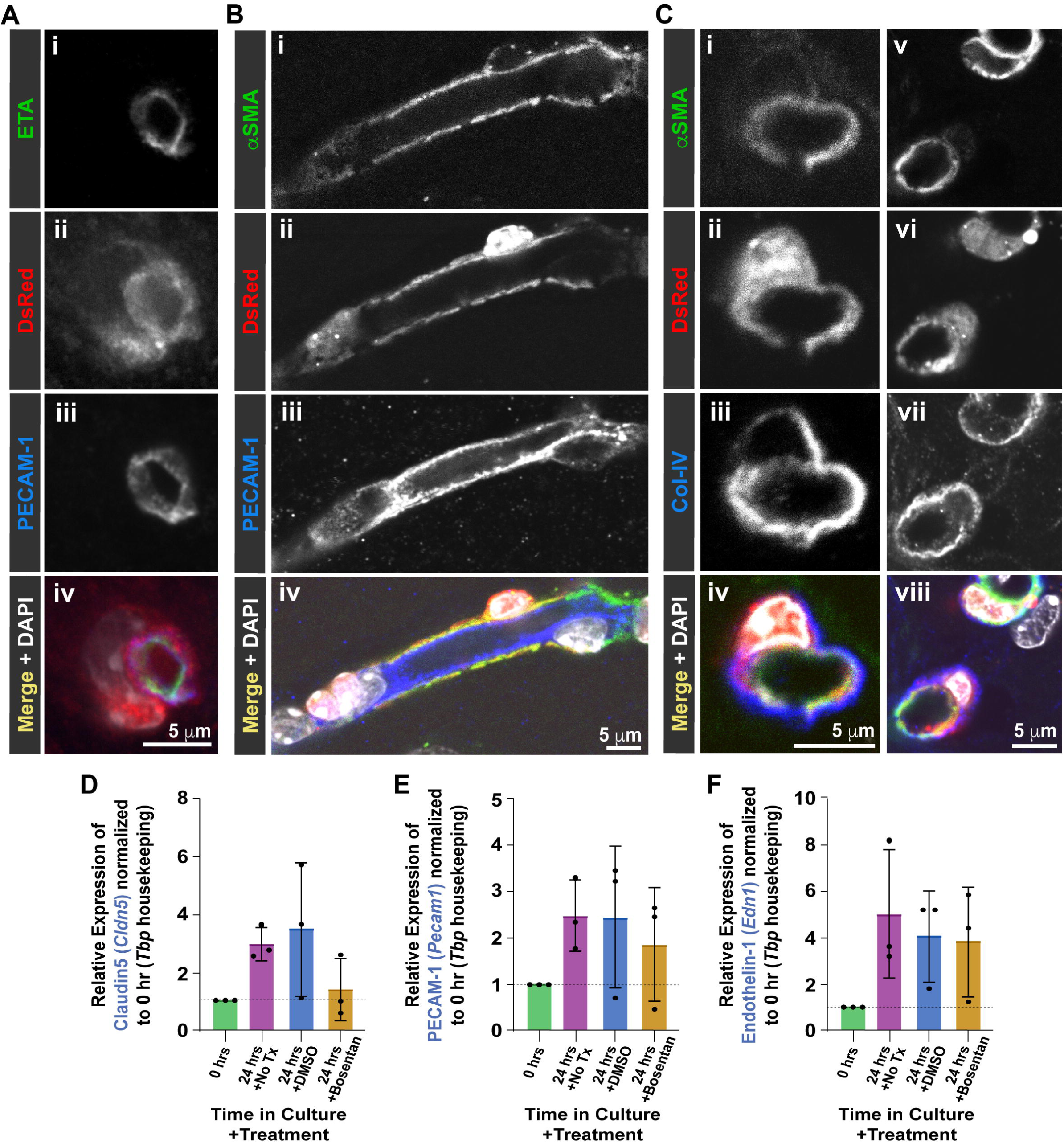
A PC Subpopulation Contained ET-1 Type A Receptors and αSMA Proteins, and Blocking ET-1 Signaling over 24-hours Limited Claudin5 Upregulation. (A) Representative images of non-cultured NG2/*Cspg4^DsRed/+^*murine brain slices with immunostained ET-1 Type A Receptors (ETA) (i, and green in iv), PCs labeled by DsRed (ii, and red in iv), and ECs labeled for PECAM-1/CD31 (iii, and blue in iv). DAPI labeled cell nuclei (white in iv). Scale bar, 5 μm. (B) Representative images of non-cultured NG2/*Cspg4^DsRed/+^*murine brain slices with immunostained α-Smooth Muscle Actin (αSMA) (i, and green in iv), PCs labeled by DsRed (ii, and red in iv), and ECs labeled for PECAM-1/CD31 (iii, and blue in iv). DAPI labeled cell nuclei (white in iv). Scale bar, 5 μm. (C) Representative images of non-cultured NG2/*Cspg4^DsRed/+^*murine brain slices with immunostained α-Smooth Muscle Actin (αSMA) (i and v, and green in iv and viii), PCs labeled by DsRed (ii and vi, and red in iv and viii), and labeled Type IV Collagen (Col-IV) (iii and vii, and blue in iv and viii). DAPI labeled cell nuclei (white in iv and viii). Scale bar, 5 μm. (D) Relative expression of *Cldn5* measured by qRT-PCR in murine brain slices cultured for 24 hours without any treatment (purple bars), with DMSO control vehicle treatment (blue bars), and with Bosentan treatment (55mM, gold bars), normalized pairwise to 0-hour controls (green bars). *Tbp* served as the housekeeping gene. Filled bars represent averages with error bars as standard deviation. Individual data points are shown for each biological replicate (n=3), and dotted gray lines indicate “1” for referencing to 0-hour control samples. (E) Relative expression of *Pecam1* measured by qRT-PCR in murine brain slices cultured for 24 hours without any treatment (purple bars), with DMSO control vehicle treatment (blue bars), and with Bosentan treatment (55mM, gold bars), normalized pairwise to 0-hour controls (green bars). *Tbp* served as the housekeeping gene. Filled bars represent averages with error bars as standard deviation. Individual data points are shown for each biological replicate (n=3), and dotted gray lines indicate “1” for referencing to 0- hour control samples. (F) Relative expression of *Edn1* measured by qRT-PCR in murine brain slices cultured for 24 hours without any treatment (purple bars), with DMSO control vehicle treatment (blue bars), and with Bosentan treatment (55mM, gold bars), normalized pairwise to 0-hour controls (green bars). *Tbp* served as the housekeeping gene. Filled bars represent averages with error bars as standard deviation. Individual data points are shown for each biological replicate (n=3), and dotted gray lines indicate “1” for referencing to 0-hour control samples.

ET-1 is well known as a potent stimulator of vSMC contraction^62^, and it is emerging as an important signaling molecule for microvascular PCs as well^63^. Recent studies have suggested that PCs, or perhaps a subpopulation within the arteriole-to-capillary transition zone, are capable of narrowing vessel diameters via their contraction^27–37^, though not without controversy^64^. ET-1 has also been implicated as a driver of PC contractility^42–44^. Within this line of investigation, questions regarding PC expression of contractile machinery proteins have also emerged, with observations supporting and refuting this conclusion^65–68^. To explore the notion of PC subtype contractility and potentially integrate this idea into our working model, we immunostained non-cultured NG2*/Cspg4^DsRed/+^* murine brain slices for α-smooth muscle actin (αSMA/*Acta2*). As before, we co-stained for PECAM-1 to mark ECs and DAPI to label cell nuclei. Vascular SMCs in arteriole-sized vessels generated a robust αSMA signal that was easily detected by confocal microscopy (data not shown), while αSMA signals along microvessels required focusing on regions where imaging parameters could be adjusted to an appropriate sensitivity. In these regions, we were able to visualize the αSMA label coinciding with DsRed signals in nuclei and cellular processes extended along PECAM-1-positive capillary ECs (Figure 6B). In a complementary set of brain sections, we replaced the PECAM-1 label with a stain for Type IV Collagen (Col-IV), an abundant ECM component within the vascular basement membrane. This approach allowed us to assess the spatial distribution of the DsRed and αSMA signals with another vascular-related molecule. We observed numerous capillary-sized vessels containing cells with overlapping DsRed and αSMA signals within an outer layer of Co-IV (Figure 6C). These observations are consistent with the notion that a subpopulation of PCs contain molecules facilitating their contractility and thus their ability to constrict vessels and reduce their diameters, consistent with their proposed response to ET-1 activation^42–44^.

Building upon these observations and exploring potential mechanistic connections within our working model, we next tested the hypothesis that elevated *Edn1* transcripts and associated ET-1 activity might be influencing the increase in *Cldn5* transcripts. Our rationale for this link was based on the idea that, in cultured brain slices lacking blood flow, ET-1 stimulation of contractile PCs may constrict capillaries^42^ such that associated ECs undergo an upregulation of *Cldn5* and tight junction remodeling. To test this experimentally, we utilized the pharmacological agent Bosentan, a known inhibitor if ET-1 signaling via both Type A and B ET-1 receptors^69,70^. We applied this drug at 55mM to our brain slice culture model for 24-hours alongside sets of slices cultured without any treatment (i.e. no Tx control) and in the presence of the drug diluent only (i.e. DMSO vehicle control). We compared *Cldn5* transcript levels from microvessels of non-cultured, 0-hour slices with cultured slices from these three experimental groups (24-hours with: 1—no Tx, 2—DMSO, or 3—Bosentan). The no treatment and DMSO groups showed increases in *Cldn5* transcripts relative to non-cultured slices, with moderate variability noted (Figure 6D). The Bosentan group however displayed an average level of *Cldn5* expression that mirrored the 0-hour group, again with some variability, but consistent with a potential dampening of Cldn5 upregulation in the presence of this ET-1 inhibitor (Figure 6D). We next asked if the increase in *Pecam1* transcript levels during 24-hour slice culture were also affected by ET-1 suppression, suggesting a broader effect on EC junctions. *Pecam1* was upregulated within all 3 groups relative to non-cultured samples (Figure 6E), suggesting that reduced ET-1 signaling activity in this context did not influence the regulation of PECAM-1-based EC junctions. Lastly, we assessed mRNA transcripts for ET-1 (*Edn1*) to ensure that none of the treatment groups experienced an unexpected decrease, confounding our interpretations. We found that all 3 groups contained elevated levels of *Edn1* transcripts relative to the 0-hour group (Figure 6F). On aggregate, these observations appear to be in good agreement with the concept that, within this specific experimental model of extended brain slice culture in aCSF, increases in *Cldn5* transcripts can be limited by blocking ET-1 activity with the pharmacological inhibitor Bosentan, suggesting a potential mechanistic link.

## DISCUSSION

Blood circulating throughout the vascular system distributes oxygen and nutrients, removes waste and cellular byproducts, and transports cells and hormonal signals throughout the body^2^. Blood also provides important mechanical cues for maintaining the structural integrity of the vasculature^1^. Perturbation or loss of these biophysical inputs can induce structural changes within the blood vessel wall that can have deleterious consequences^3–9^. In the current study, we analyzed the microvasculature from non-cultured murine brain slices and those cultured in oxygenated aCSF for 12- and 24-hours to better understand how cerebral ECs and PCs respond to blood flow cessation. Consistent with other scenarios of altered hemodynamics^8,9^, molecular mediators of inflammation were upregulated in cultured slices, particularly those associated leukocyte recruitment and infiltration. Transcriptional signatures of ECM remodeling and cell-ECM interactions were also elevated within cultured cortical microvessels lacking blood flow. These changes were likely related to the observed reduction in PC coverage of cerebral capillaries. We presumed that these responses were indicative of BBB breakdown and degradation of the associated endothelium. To our surprise, we found that: (i) mRNA transcripts for EC junctions increased after 24-hours in culture, (ii) protein levels for junction molecules remained relatively stable, and (iii) Claudin5-based tight junctions were structurally rearranged. We also noted a reduction in the outer diameters of numerous, but not all, capillaries in cultured brain slices, suggesting that a subset of microvessels may have been constricted by associated PCs. Consistent with this interpretation, we found molecular hallmarks of vasoconstrictor ET-1 upregulation within brain slices cultured over time in aCSF. Furthermore, we detected ET-1 receptors and contractile machinery proteins (i.e. αSMA) within a subpopulation of cerebral PCs, supporting a potential role in capillary constriction. We then found that suppressing ET-1 activity in this no-flow brain slice model prevented an increase in Claudin5/*Cldn5* regulation. Taken together, these observations suggest that, in this experimental murine brain slice model of cerebral blood flow cessation, a subpopulation of PCs contribute to capillary wall remodeling by reducing ECM components within the vBM and detaching from the wall altogether. Our data further suggest that a potential mechanistic link may exist between ET-1 activity and Claudin5 regulation during scenarios of blood flow cessation, mediated in part via constriction of the cerebral microvasculature by a subset of PCs.

Capillary perfusion can be reduced or lost altogether in several clinically relevant pathologies. In coronary or peripheral artery disease, for example, atherosclerotic lesions can limit blood flow into the microcirculation as they expand into the lumen of upstream feeder arteries^71,72^. Unstable plaques can also rupture, giving rise to clots that suddenly impede blood flow to downstream tissues, as seen in myocardial infarction and ischemic stroke^73^. Clinical management of these pathologies focuses primarily on blood flow restoration, but it has been observed that, despite successful removal of a blockage, complete reperfusion of downstream tissues can be limited^10,74^. This scenario has been described as the “no re-flow” phenomena^75,76^. Microvascular PCs have been implicated in preventing the return of blood flow into capillaries via vessel constriction and subsequent cell death i.e. PC *rigor mortis*^38–41^. Our data suggest that an additional confounding factor in restoring tissue perfusion might be that PC-induced capillary collapse may cause EC junctions to remodel and limit vessel lumen re-expansion. These EC junctional rearrangements appear to be sensitive to ET-1 upregulation, which is consistent with previous studies observing ET-1-driven PC constriction^42^. Exposure to abnormally high levels of oxygen (i.e. hyperoxia) has been linked with ET-1 upregulation but not directly with PC constriction ^32^. Prior studies have also described hypoxia as an inducer of ET-1 release^45^, further highlighting the complexities of these mechanistic relationships. Our data suggests that the rapid change in hemodynamics may also be a significant driver of these responses including ET-1 upregulation, as reported previously^46^. Furthermore, we cannot rule out the potential contributions of directly exposing brain slices to a relatively high level of sugar in the aCSF formulation^77^. Thus, further work will be necessary to dissect the degree to which these environmental inputs (i.e. altered hemodynamics, oxygen tension, glucose levels) are integrated to induce the observed responses from microvascular ECs and PCs in the cerebrovasculature and within capillary beds throughout the body.

Pericyte detachment from the capillary wall is a common feature in numerous pathologies involving microvascular dysfunction. Diabetes mellitus for example involves PC loss from microvessels in the retina^15,21^, among other tissues^78^. This phenomenon is a hallmark of various vascular-related complications including those that involve sudden or progressive changes in hemodynamics and tissue perfusion^79^. A specific subpopulation of PCs may be more susceptible to capillary detachment, particularly those characterized as “thin-strand PCs” (tsPCs) that extend thin cellular processes along the capillary wall^22,36,80^. We have previously observed tsPCs as they retract these extensions towards their soma, seemingly in preparation to migrate outward from the microvasculature^81^. The ECM ensheathing tsPCs must be degraded to allow for “escape”, which includes the release of MMP9 around the cell body^23,25^. In the present study, we also observed a notable increase in *Mmp9* expression at 12- and 24-hours after blood flow cessation in murine brain slices cultured within aCSF, consistent with these previous studies. Interestingly, we found a rapid and sustained decline in vitronectin mRNA and protein, which agrees with previous literature implicating this ECM component as an important element for PC integration into the vessel wall^13,14^. It appears that cerebral capillary ECs may have upregulated Integrin-α5 after 24-hours in culture as a compensatory response to reduced vitronectin levels, given that ECs utilize this integrin to bind PC-derived vitronectin^14^. However, this important ECM receptor has also been implicated as a blood flow sensor in conjunction with Vascular Endothelial Growth Factor Receptor-2 (VEGFR2) and PECAM-1, suggesting that the observed increases in *Itga5* and *Pecam1* transcripts at 24-hours may also be related to their role in flow-sensing mechanisms in capillary ECs^82^. Overall, our data align with previous studies suggesting that PC detachment from the capillary wall is preceded by distinct changes in the ECM composition of the vascular basement membrane, which may be sensitive to altered hemodynamics within the microcirculation.

As with all scientific inquiry, we must consider important limitations within the current study alongside our interpretations of the data and the associated literature. Herein, we assessed PC coverage by visualizing DsRed signals generated by the NG2*/Cspg4^DsRed/+^* reporter construct as well as immunostaining for PDGFRβ. There is a possibility that the reduction of these signals along cultured brain microvessels reflects a reduction in marker expression and was not solely due to PC detachment from the capillary wall. Follow-on studies could include a more comprehensive transcriptional profiling of cerebral PCs to determine if the loss of blood flow and these specific culture conditions affected mechanisms maintaining PC identity and their molecular expression pattern. Such an approach could also be coupled with developing rigorous methods for quantifying capillary diameters across a given brain section. Spatial mapping of these measures might reveal correlations between microvessel diameter changes, EC junction morphology, and the expression profiles of PC subpopulations. Another opportunity for advancing this line of inquiry is exploring the potential involvement of Zonula Occludens-1 (ZO-1/Tjp1) or Vascular-Endothelial Cadherin (VE-Cadherin/*Cdh5*), which is another important EC junction molecule that is flow-regulated^83^ and may be affected in a similar manner as Claudin5. Lastly, orthogonal imaging approaches could offer new insight into responses observed within these no-flow brain slices cultured in aCSF. For instance, time-lapse imaging of fluorescently labeled PCs and ECs, as we have done previously^37,81,84–86^, could capture the temporal dynamics of PC subtypes detaching from capillaries and/or constricting microvessels over the 12- and 24-hour time periods. This modality could be applied to tissues with and without simulated perfusion via large-vessel catheterization and a flow pump system, as we have begun developing for other tissues including the intestine^50^. Serial block face-scanning electron microscopy (SBF-SEM) could also be applied to non-cultured^80^ and cultured samples to determine on an ultrastructural level: (i) if constricted capillaries are indeed fully collapsed, and (ii) how the cellular and ECM composition within cerebral microvessels has remodeled at different time points. While this is certainly not an exhaustive list of potential limitations with the current study, these are areas for further exploration and opportunities for new insights within follow-on efforts. Overall, the observations presented herein contribute to our collective understanding of how PCs and ECs within cerebral capillaries may adapt to environmental changes.

Given that a primary feature of the brain slice model is the sudden loss of blood flow, we focused our initial interpretations on the impact of altered hemodynamics; however, additional aspects of our experimental set up (e.g. oxygen tension, glucose levels) might also be integrated in the responses observed^21,32,45,77^. We attempted to address potential effects of the altered glucose environment, though a shorter time-course for diet-induced hyperglycemia might be more appropriate, or perhaps direct exposure to elevated glucose is needed. Nevertheless, while more work will be needed to more precisely dissect the inputs driving these cellular and molecular outcomes, these data align with previous literature suggesting that PC subpopulations and the cerebral endothelium undergo coordinated responses to disrupted environmental conditions^87,88^. ECM remodeling and the loss of PCs from brain capillaries during no-flow conditions might be an important consideration for clinical interventions aimed at restoring flow^38–41,75,76^, as microvessel stability may be altered and lead to impaired perfusion and tissue exchange and even rupture. Furthermore, if a PC subset constricts capillaries and leads to EC junction remodeling, suggested by the alignment of our data with previous studies^11,12,27–37^, then therapeutic efforts to restore perfusion may be limited if these microvessels are structurally unable to regain a patent lumen. Additional studies will be necessary to more fully elucidate these mechanisms and determine molecular targets for improving therapeutic outcomes.

## Supplemental Information

Supplemental information can be found online and includes data from a model of diet-induced hyperglycemia, original western blot images, additional confocal images, a graphical summary, and data from publicly available databases associated with published studies.

## Acknowledgements.

We would like to thank all the members of the Chappell lab for their support and valued assistance on this project, both materially and intellectually. This work was supported in part by funding from the National Institutes of Health (R01HL146596 to J.C.C., F31HL168946 to H.A., R01HL168559 to J.P., and F31HL174075 to M.W.) and the American Heart Association (AHA, 19TPA34910121 to J.C.C.).

## Author Contributions

H.A. and J.C.C. conceptualized and optimized methodology. H.A., M.A., and M.W. performed experiments. H.A., M.W., and J.C.C. analyzed datasets. H.A. and J.C.C. wrote the manuscript. H.A. and M.W. provided critical data for the manuscript. H.A., M.W., J.P., and J.C.C. acquired financial support. All authors have reviewed and approved the final version of this manuscript.

## Inclusion and Diversity

We support inclusive, diverse, and equitable conduct of research.

## Data Availability

The datasets generated during and/or analyzed during the current study are included in the manuscript, and all supporting data are available from the corresponding author on reasonable request.

## Abbreviations

EC: endothelial cell
PC: pericyte
BBB: blood-brain barrier
ECM: extracellular matrix
vBM: vascular basement membrane
tsPC: thin-strand pericyte
MMP: matrix metalloproteinase
ePC: ensheathing pericyte
mPC: mesh pericyte
ET-1: Endothelin-1
aCSF: artificial cerebrospinal fluid
Vtn: vitronectin
αSMA: α-smooth muscle actin
vSMC: vascular smooth muscle cell.

