## Supplemental Material for "Pericyte and Endothelial Cell Responses within Murine Cerebral Capillaries After Blood Flow Cessation"

### SUPPLEMENTAL INFORMATION

**Supplemental Figure 1. Diet-induced Hyperglycemia Does Not Induce Similar Transcription Changes in Murine Cerebral Microvessels as Observed in the Brain Slice Culture Model.** (A) Fasted blood sugar levels (mg/dL) from WT mice fed a control diet (blue bar) or a high fat-high sugar (HF/HS) diet (gold bar) for 16 weeks. Bars represent averages with standard deviations, and biological replicates (n=6) are shown as individual data points. * denotes $p \leq 0.05$ by unpaired Student T-test comparison. (B) Body weight (g) of WT mice fed a control diet (blue bar) or a high fat-high sugar (HF/HS) diet (gold bar) for 16 weeks. Bars represent averages with standard deviations, and biological replicates (n=6) are shown as individual data points. * denotes $p \leq 0.05$ by unpaired Student T-test comparison. Relative expression of (C) Vitronectin (*Vtn*), (D) PECAM-1 (*Pecam1*), (E) Claudin5 (*Cldn5*), and (F) Endothelin-1 (*Edn1*) measured by qRT-PCR in murine brain microvessels from WT mice fed a control diet (blue bar) or a high fat-high sugar (HF/HS) diet (gold bar) for 16 weeks. *Tbp* served as the housekeeping gene. Filled bars represent averages with error bars as standard deviation. Individual data points are shown for 2 pooled biological replicates (n=3 replicates from 6 subjects), and dotted gray lines indicate "1" for referencing to Control Diet samples. Unpaired Student T-test comparisons did not detect any significant differences in gene expression.

**Supplemental Figures 2-4.** Original western blots in support of Figures 2 and 4.

**Supplemental Figure 5. Claudin5 Morphological Rearrangement Visualized in Cerebral Capillaries within Murine Brain Slices Cultured in aCSF for 24-hours.** Representative images of WT murine brain slices following 0-hours (A) and 24-hours (B) in culture with ECs labeled for PECAM-1/CD31 (for both A and B: i, and blue in iv) and for Claudin5 (for both A and B: ii, and green in iv), and PCs labeled for PDGFR$\beta$ (for both A and B: iii, and red in iv). DAPI labeled cell nuclei (for both A and B: white in iv). Scale bars, 5 $\mu$m. For both A and B, a single focal plane from a z-stack is shown in i-iv (x-y axis shown in lower left corner), and a reconstructed orthogonal view at the location indicated by the white dotted line is shown in v-viii (z-y axis shown in the lower left corner).

**Supplemental Figures 6 and 7.** Original western blots and additional biological replicates in support of Figure 5.

**Supplemental Figure 8. Summary Schematic of our Working Hypothesis.** This conceptual framework posits that (i) cerebral capillary ECs (blue) release ET-1 (dark blue) during slice culture, (ii) PCs (red), or perhaps a subset, are responsive to increased ET-1 levels via cell surface receptors (dark red), and (iii) these signals stimulate this PC subpopulation to constrict and collapse a portion of capillaries, leading to the observed changes in Claudin5 (green) transcription and morphology.

**Supplemental Figure 9. Vascular Smooth Muscle Cells Harbor ET-1 Type A Receptors in Non-Cultured Murine Brain Slices.** Providing a positive control for ETA antibody labeling in Figure 6, representative images of non-cultured NG2/*Cspg4*$^{DsRed/+}$ murine brain slices (i.e. 0-hours in culture) with immunostained ET-1 Type A Receptors (ETA) (i, and green in iv), vascular smooth muscle cells (vSMCs) labeled by DsRed (ii, and red in iv), and ECs labeled for PECAM-1/CD31 (iii, and blue in iv). DAPI labeled cell nuclei (white in iv). Scale bar, 20 μm. Tan arrows in i-iv indicate ETA-positive vSMCs also positive for DsRed and adjacent to ECs within a larger-diameter cerebral artery. A single focal plane from a z-stack is shown in i-iv.

**Supplemental Figure 10. Single Cell RNA-sequencing Databases Provide Measurements Aligning with the Presence of *Ednra* (ET-1 Type A Receptor, ETA) Transcripts in Presumptive PC populations from Various Murine Tissue Sources.** (A) Plot of normalized *Ednra* expression in main cluster and subclusters consistent with PC precursor identity (see Payne et al. *ATVB* 2022) from Mouse Organogenesis Cell Atlas (MOCA, Shendure & Trapnell Labs – Cao et al. *Nature* 2019). See publication for complete experimental details. (B) Plot of normalized *Ednra* expression in murine brain samples from single cell RNA Sequencing of WT Murine Brain, associated with the following publications: Vanlandewijck, He et al. *Nature* 2018, He, Vanlandewijck et al. *Scientific Data*, 2018. See publications for complete experimental details. (C) Plot of normalized *Ednra* expression from single cell RNA Sequencing of WT Murine Brain, specifically from the visual cortex, associated with the following publication: Hrvatin, Hochbaum, Nagy et al. *Nat Neurosci* 2018. See publication for complete experimental details.

**Supplemental Figure 1.**

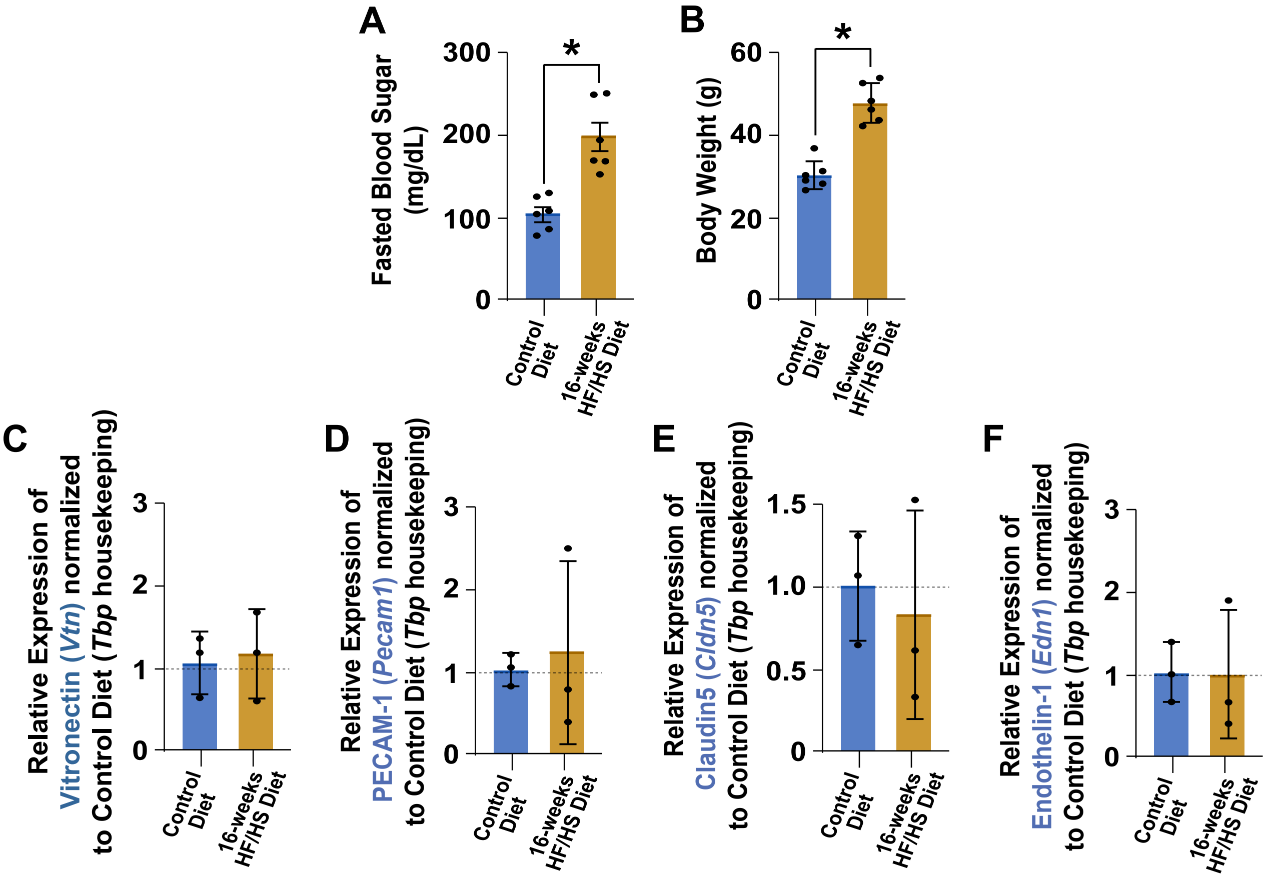

**Supplemental Figure 2.**

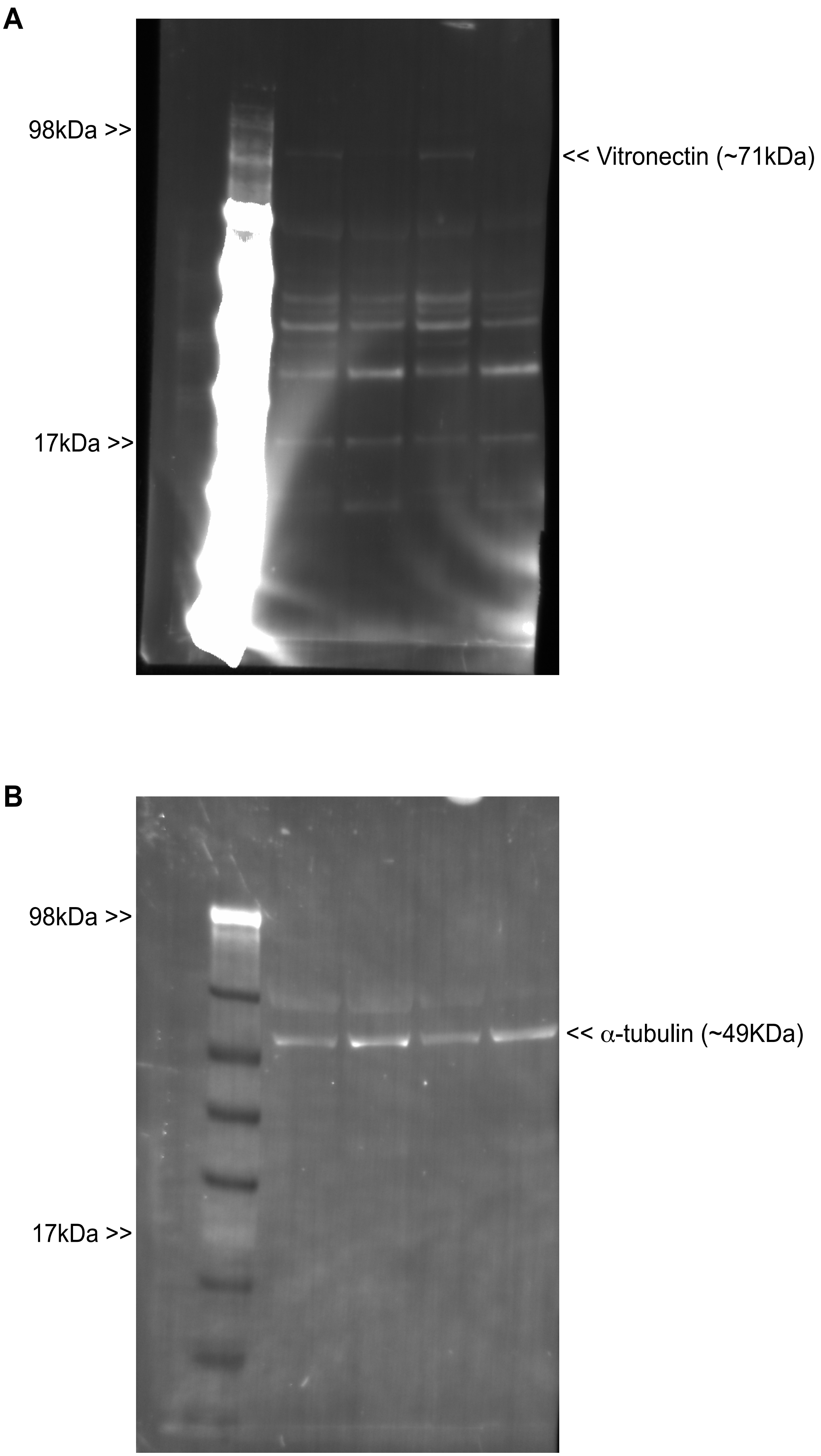

**Supplemental Figure 3.**

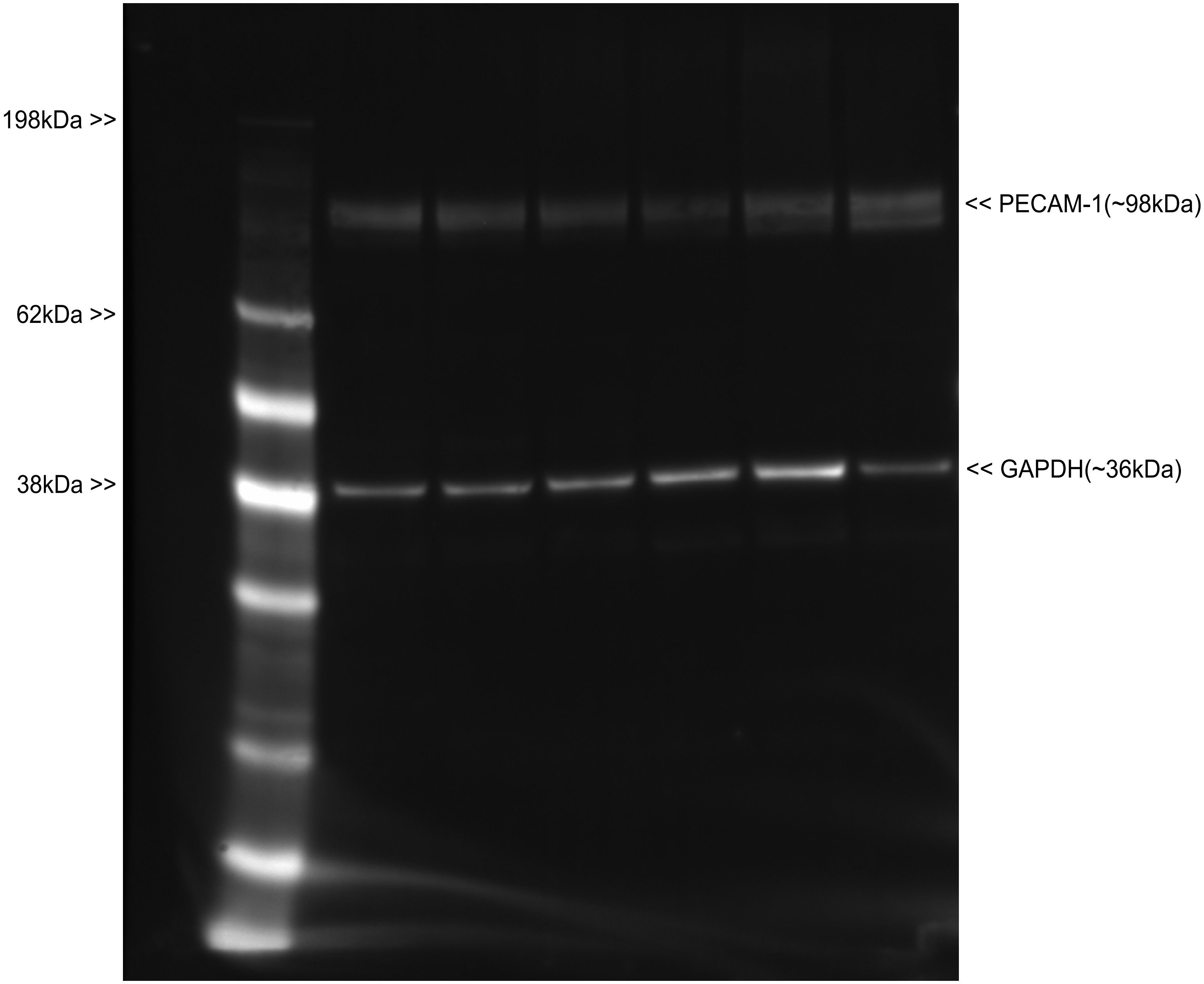

**Supplemental Figure 4.**

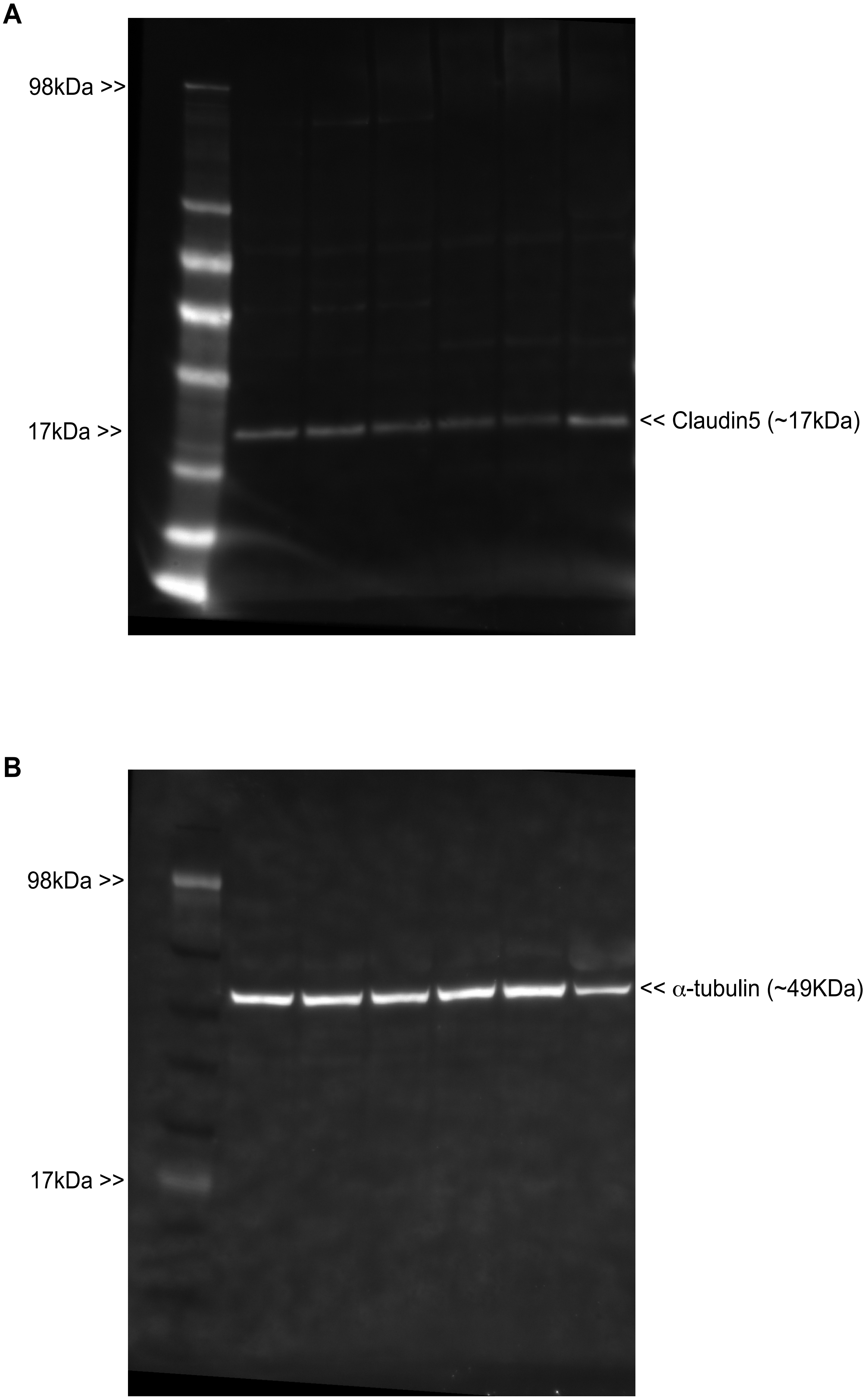

**Supplemental Figure 5.**

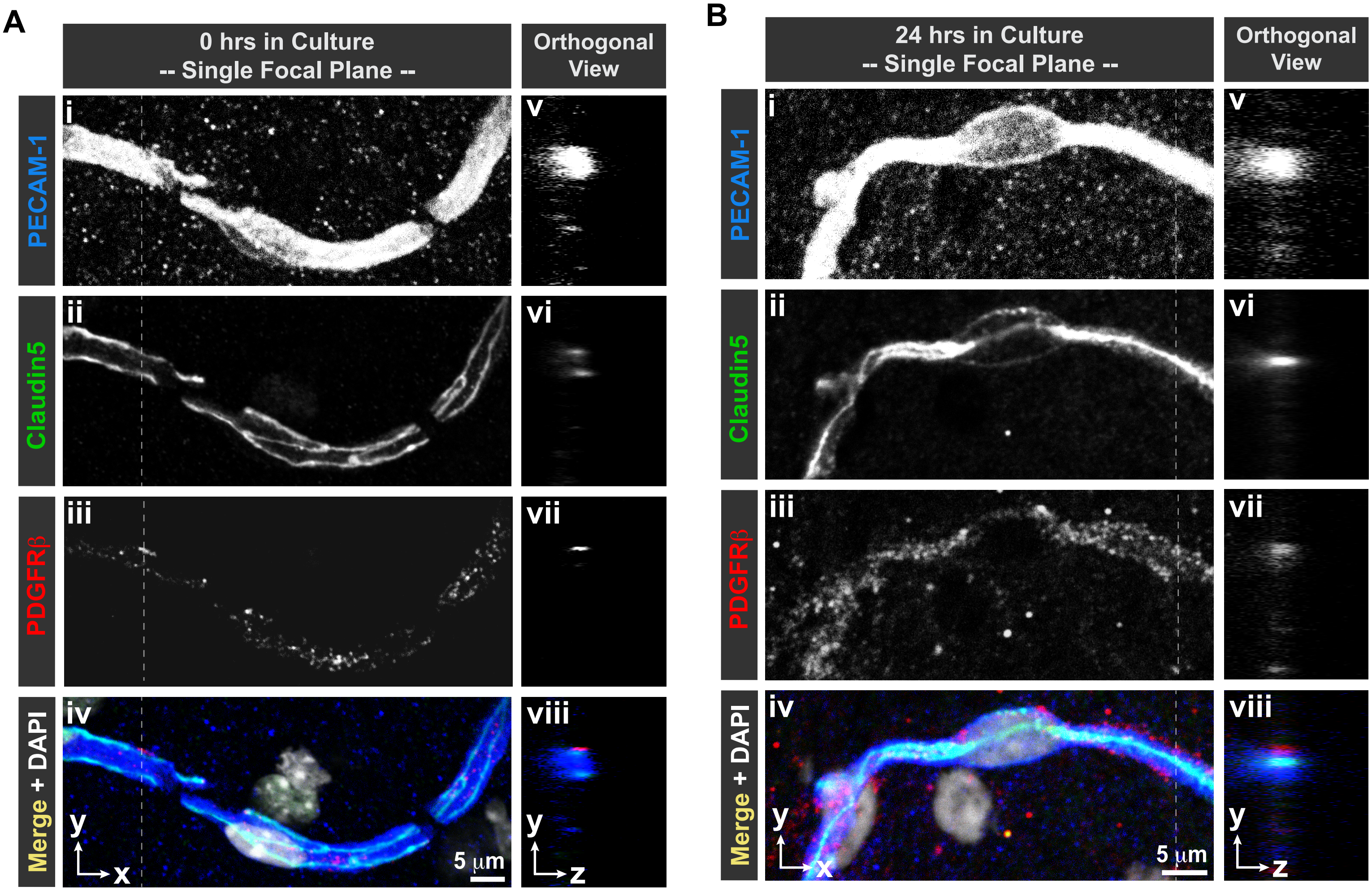

**Supplemental Figure 6.**

A

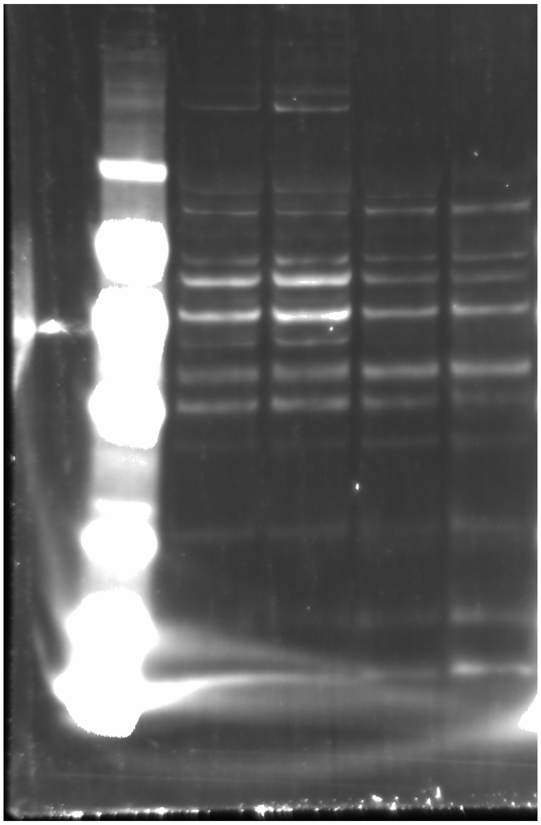


B

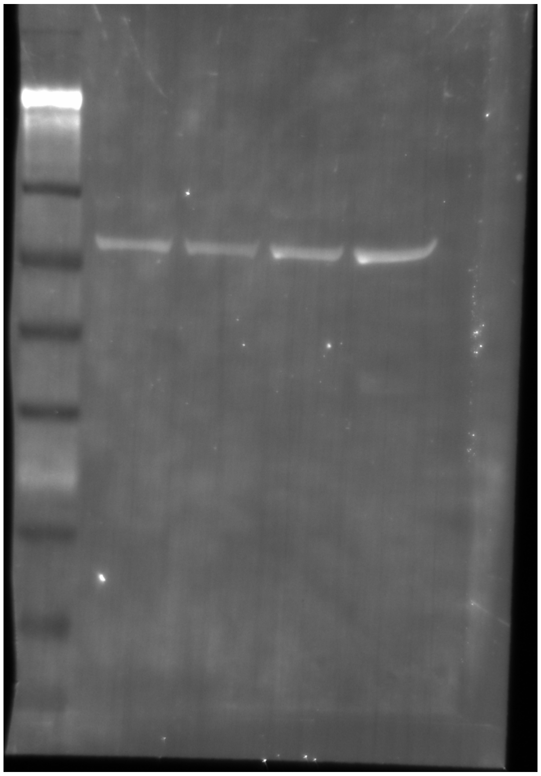

**Supplemental Figure 7.**

A

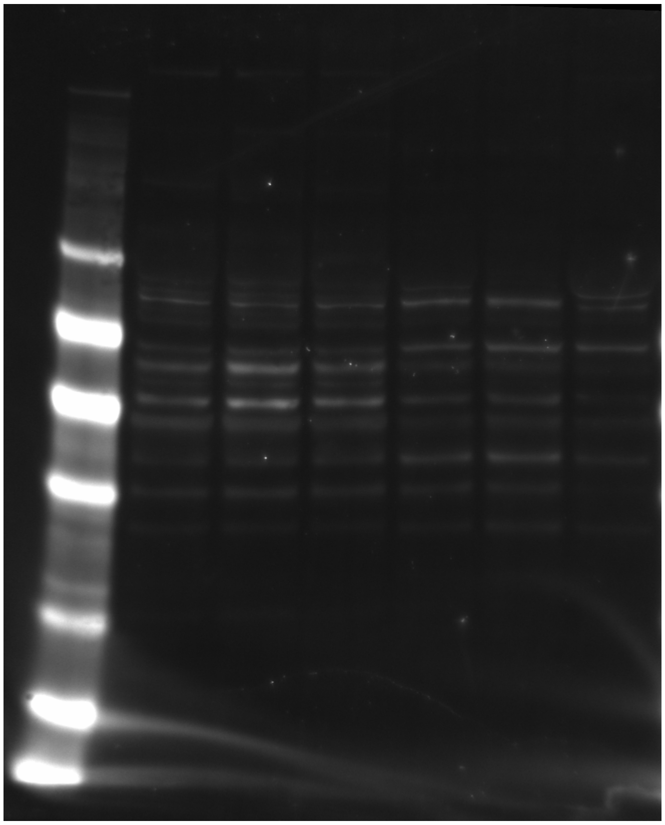


B

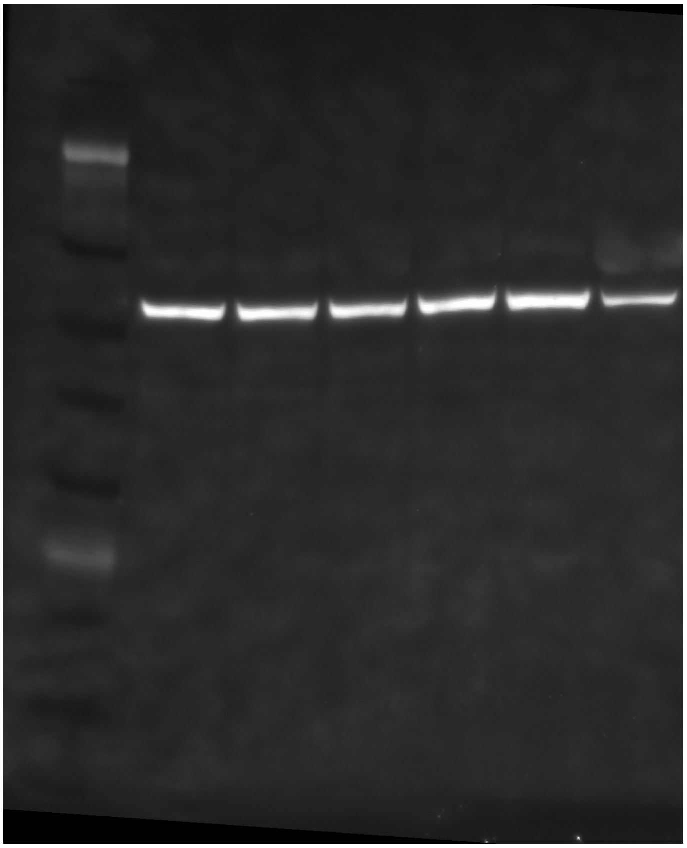

Supplemental Figure 8.

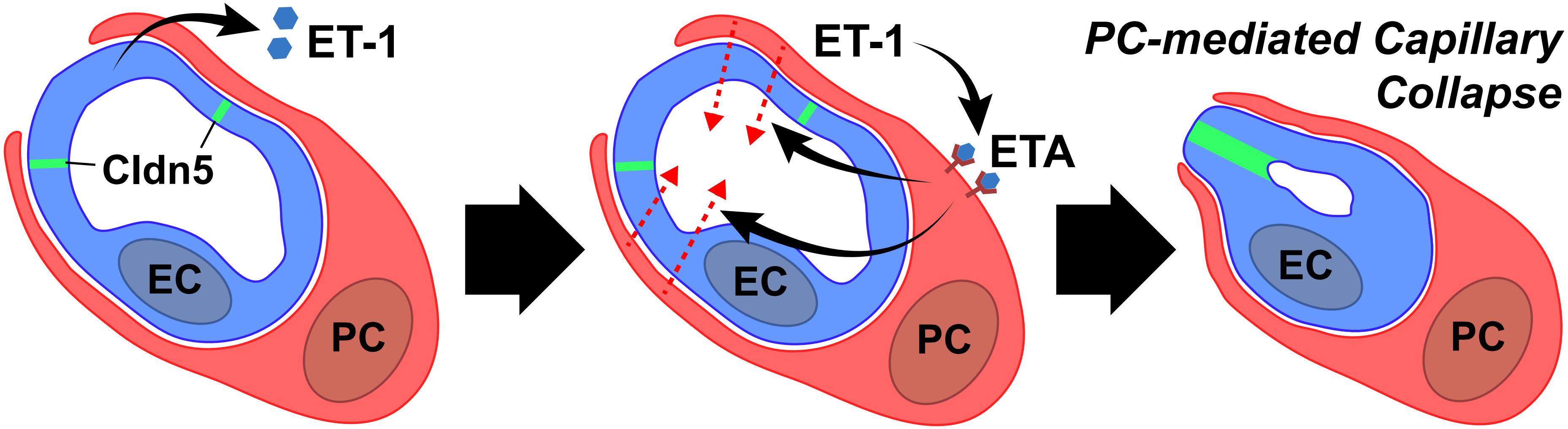

**Supplemental Figure 9.**

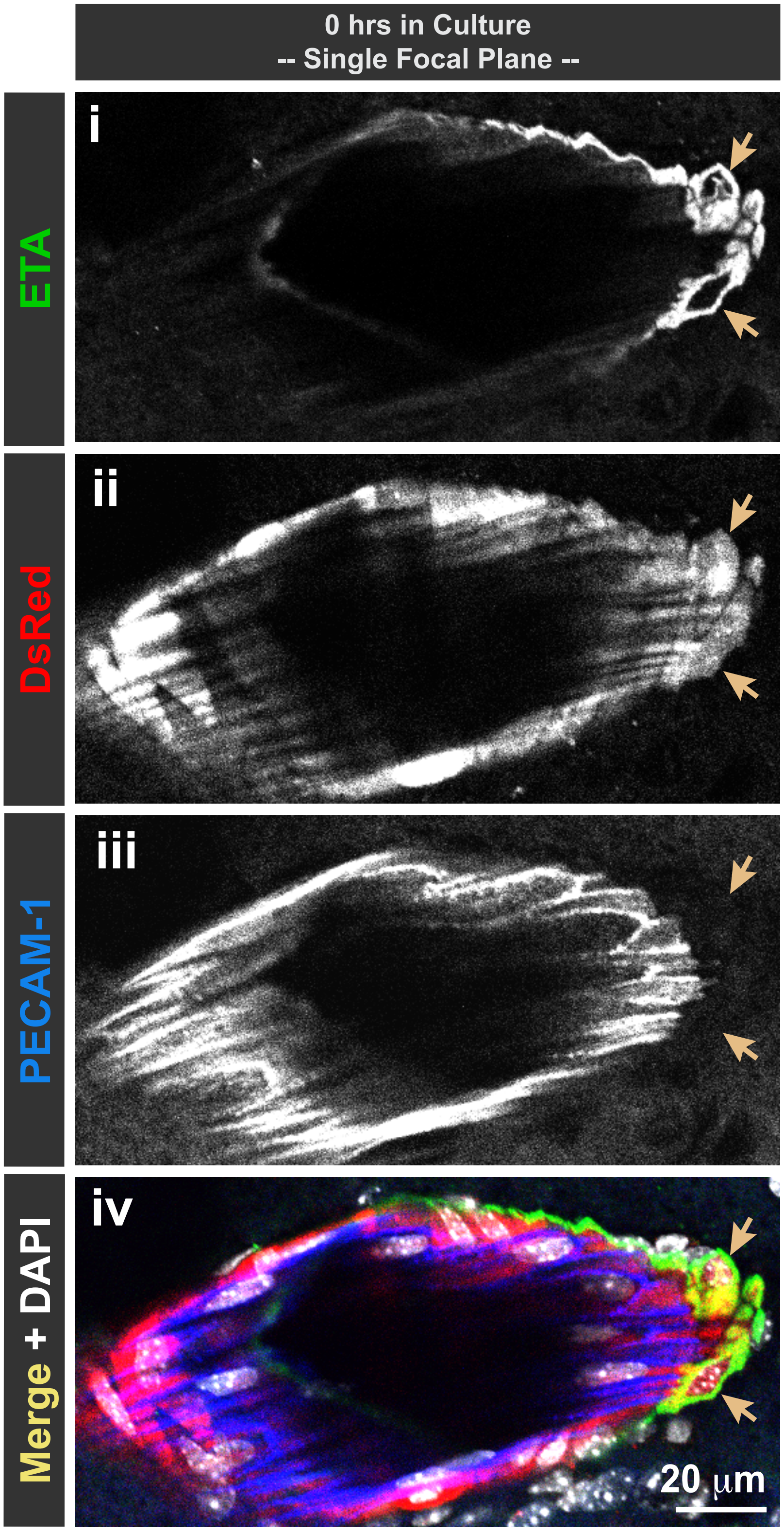

### Supplemental Figure 10.

**A**

**Plot of normalized expression in main cluster and subclusters, from Mouse Organogenesis Cell Atlas (MOCA), Shendure & Trapnell Lab**

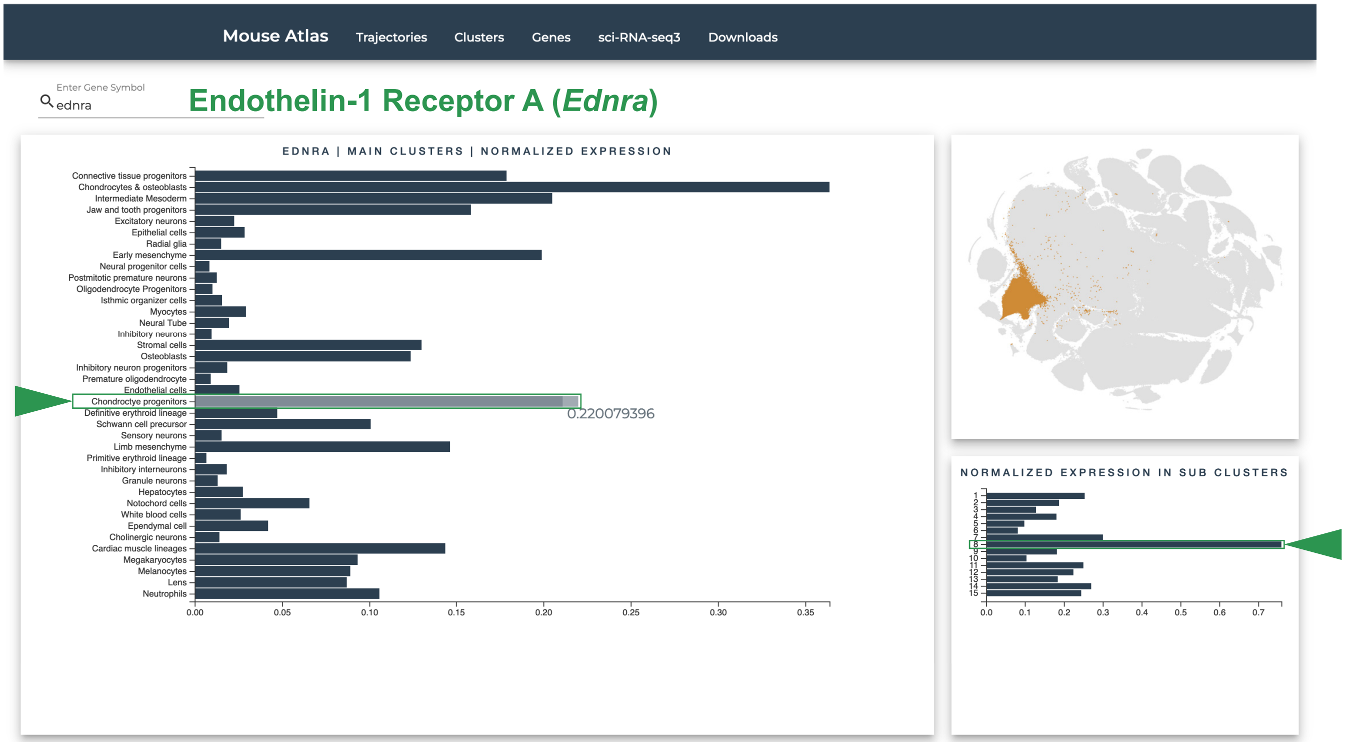


**B** **Endothelin-1 Receptor A (*Ednra*)**

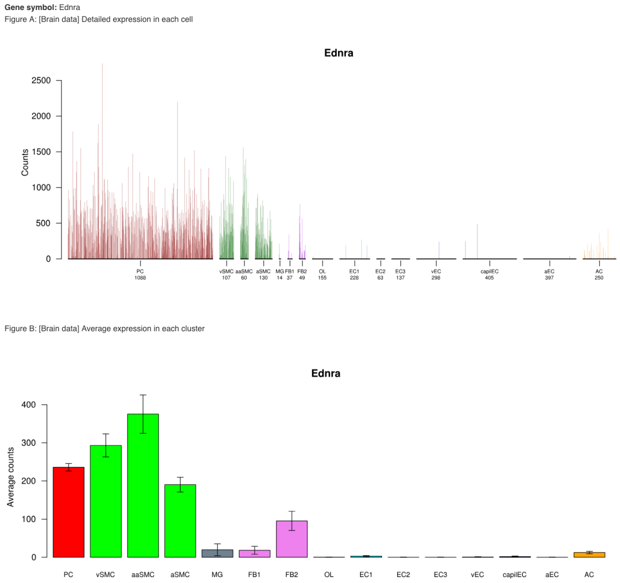


***Abbreviations:***

**[Brain data]: PC - Pericytes; SMC - Smooth muscle cells; MG - Microglia; FB - Vascular fibroblast-like cells; OL - Oligodendrocytes; EC - Endothelial cells; AC - Astrocytes; v - venous; capil - capillary; a - arterial; aa - arteriolar; 1,2,3- subtypes.**

**Plot of normalized expression from single cell RNA Sequencing of WT Murine Brain, http://betsholtzlab.org/VascularSingleCells/database.html associated with the following publications: Vanlandewijck, He et al. Nature 2018, He, Vanlandewijck et al. Scientific Data, 2018**

**C** **Endothelin-1 Receptor A (*Ednra*)**

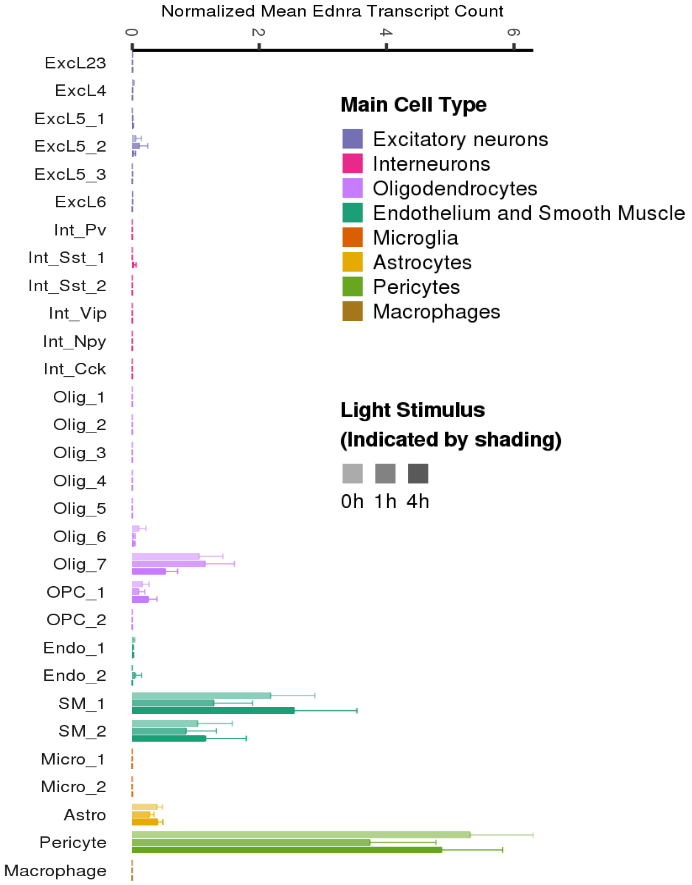


**Plot of normalized expression from single cell RNA Sequencing of WT Murine Brain, specifically visual cortex. https://greenberg.hms.harvard.edu/gene-database/ associated with the following publications: Hrvatin, Hochbaum, Nagy et al. *Nat Neurosci* 2018. See publication and website for complete experimental details.**
